# Advancing Divide-and-Conquer Phylogeny Estimation using Robinson-Foulds Supertrees

**DOI:** 10.1101/2020.05.16.099895

**Authors:** Xilin Yu, Thien Le, Sarah A. Christensen, Erin K. Molloy, Tandy Warnow

## Abstract

One of the Grand Challenges in Science is the construction of the *Tree of Life*, an evolutionary tree containing several million species, spanning all life on earth. However, the construction of the Tree of Life is enormously computationally challenging, as all the current most accurate methods are either heuristics for **NP**-hard optimization problems or Bayesian MCMC methods that sample from tree space. One of the most promising approaches for improving scalability and accuracy for phylogeny estimation uses divide-and-conquer: a set of species is divided into overlapping subsets, trees are constructed on the subsets, and then merged together using a “supertree method”. Here, we present Exact-RFS-2, the first polynomial-time algorithm to find an optimal supertree of two trees, using the Robinson-Foulds Supertree (RFS) criterion (a major approach in supertree estimation that is related to maximum likelihood supertrees), and we prove that finding the RFS of three input trees is **NP**-hard. We also present GreedyRFS (a greedy heuristic that operates by repeatedly using Exact-RFS-2 on pairs of trees, until all the trees are merged into a single supertree). We evaluate Exact-RFS-2 and GreedyRFS, and show that they have better accuracy than the current leading heuristic for RFS. Exact-RFS-2 and GreedyRFS are available in open source form on Github at github.com/yuxilin51/GreedyRFS.

## 1 Introduction

Supertree construction (i.e., the combination of a collection of trees, each on a potentially different subset of the species, into a tree on the full set of species) is a natural algorithmic problem that has important applications to computational biology; see [7] for a 2004 book on the subject and [9, 10, 14, 16–18, 27, 40] for some of the recent papers on this subject. Supertree methods are particularly important for large-scale phylogeny estimation, where it can be used as a final step in a divide-and-conquer pipeline [48]: the species set is divided into two or more overlapping subsets, unrooted leaf-labelled trees are constructed (possibly recursively) on each subset, and then these subset trees are combined into a tree on the full dataset, using the selected supertree method. Furthermore, provided that optimal supertrees are computed, divide-and-conquer pipelines can be provably *statistically consistent* under stochastic models of evolution: i.e., as the amount of input data (e.g., sequence lengths when estimating gene trees, or number of gene trees when estimating species trees) increases, the probability that the true tree is returned converges to 1 [26, 47].

Unfortunately, the most accurate supertree methods are typically heuristics for **NP**-hard optimization problems [4, 16, 27, 29, 31, 35, 36, 40], and so are computationally intensive. However, divide-and-conquer strategies, especially recursive ones, may only need to apply supertree methods to two trees at a time, and hence the computational complexity of supertree estimation given two trees is of interest. One optimization problem where optimal supertrees can be found on two trees is the **NP**-hard Maximum Agreement Supertree (SMAST) problem (also known as the Agreement Supertree Taxon Removal problem), which removes a minimum number of leaves so that the reduced trees have an agreement supertree [14, 17]. Similarly, the Maximum Compatible Supertree (SMCT) problem, which removes a minimum number of leaves so that the reduced trees have a compatibility supertree [5, 6], can also be solved in polynomial time on two trees (and note that SMAST and SMCT are identical when the input trees are fully resolved). Because SMAST and SMCT remove taxa, methods for these optimization problems are not *true supertree methods*, because they do not return a tree on the entire set of taxa. However, solutions to SMAST and SMCT could potentially be used as *constraints* for other supertree methods, where the deleted leaves are added into the computed SMAST or SMCT trees, so as to optimize the desired criterion.

When restricting to methods that return trees on the full set of taxa, much less seems to be understood about finding supertrees on two trees. However, if the two input trees are compatible (i.e., there is a supertree that equals or refines each tree when restricted to the respective leaf set), then finding that compatibility supertree is solvable in polynomial time, using (for example) the well known BUILD algorithm [1], but more efficient algorithms exist (e.g., [3, 6]).

Since compatibility is a strong requirement (rarely seen in biological datasets), optimization problems are more relevant. One optimization problem worth discussing is the Maximum Agreement Supertree Edge Contraction problem (which takes as input a set of rooted trees and seeks a minimum number of edges to collapse so that an agreement supertree exists). This problem is **NP**-hard, but the decision problem can be solved in *O*((2*k*)^*p*^*kn*^2^) time when the input has *k* trees and *p* is the allowed number of number of edges to be collapsed [14]. Note that this analysis means that their algorithm for AST-EC may be exponential even for two trees, when the number of edges that must be collapsed is *Ω*(*n*) (e.g., imagine two caterpillar trees, where one is obtained from the other by moving the left-most leaf to the rightmost position).

In sum, while supertree methods are important and well studied, when restricted to the major optimization problems that do not remove taxa, polynomial time methods do not seem to be available, even for the special case where the input contains just two trees. This restriction has consequences for large-scale phylogeny estimation, as without good supertree methods, divide-and-conquer pipelines are not guaranteed to be statistically consistent, are not fast, and do not have good scalability [47].

In this paper we examine the well known Robinson-Foulds Supertree (RFS) problem [2], which seeks a supertree that minimizes the total Robinson-Foulds distance to the input trees. Although RFS is **NP**-hard [22], it has several desirable properties: it is closely related to maximum likelihood supertrees [38] and, as shown very recently, has good theoretical performance for species tree estimation in the presence of gene duplication and loss [25]. Because of its importance, there are several methods for RFS supertrees, including PluMiST [18], MulRF [8], and FastRFS [44]. A comparison between FastRFS and other supertree methods (MRL [27], ASTRAL, ASTRID [43], PluMiST, and MulRF) on simulated and biological supertree datasets showed that FastRFS matched or improved on the other methods with respect to topological accuracy and RFS criterion scores [44]. Hence, FastRFS is currently the leading method for the RFS optimization problem.

The main contributions of this paper are:

– We prove that RFS is solvable in *O*(*n*^2^ |*X*|) time for two trees, where *n* is the number of leaves and *X* is the number of shared leaves (Theorem 1) and **NP**-hard for three or more trees (Lemma 6).
– We present Exact-RFS-2, a polynomial time algorithm for the RFS problem when given only two source trees, and explore its performance on simulated data, both within a natural divide-and-conquer pipeline and within a greedy heuristic (Section 3). We show that Exact-RFS-2 outperforms FastRFS [44] on two trees, the current most accurate method for RFS, and that GreedyRFS is better than FastRFS for small to moderate numbers of source trees (Section 4).
– We prove that divide-and-conquer pipelines using Exact-RFS-2 are statistically consistent methods for phylogenetic tree estimation (both gene trees and species trees) under standard sequence evolution models (Theorem 2).
– We establish equivalence between RFS and some other supertree problems (Lemma 1).
– We show critical differences between RFS and SMAST/SMCT problems, that establish that methods for SMAST or SMCT cannot provably be used to constrain the search for RFS supertrees (Lemma 23).

The remainder of the paper is organized as follows. In Section 2, we provide terminology and define the optimization problems we consider. We present the Exact-RFS-2 algorithm and establish theory related to the algorithm in Section 3. Our experimental performance study is presented in Section 4, and we conclude in Section 5 with a discussion of trends and future research directions.

The details of the performance study and commands necessary to reproduce the study are omitted from the main paper but available in the appendices; the proofs are also available in [50]. All datasets used in this study are publicly available from prior studies, and the scripts and codes necessary to reproduce the study are available at http://github.com/yuxilin51/GreedyRFS.

## 2 Terminology and Problem Statements

We let [*N*] = {1, 2, *…, N*} and *𝒜* = {*T*_*i*_ | *i ∈* [*N*]} denote the input to a supertree problem, where each *T*_*i*_ is a phylogenetic tree on leaf set *L*(*T*_*i*_) = *S*_*i*_ ⊆ *S* (where *L*(*t*) denotes the leaf set of *t*) and the output is a tree *T* where *L*(*T*) is the set of all species that appear as a leaf in at least one tree in *𝒜*, which we will assume is all of *S*. We use the standard supertree terminology, and refer to the trees in *𝒜* as “source trees” and the set *𝒜* as a “profile”.

### Robinson-Foulds Supertree

Each edge *e* in a tree *T* defines a bipartition *π*_*e*_ := [*A*|*B*] of the leaf set, and each tree is defined by the set *C*(*T*) := {*π*_*e*_ | *e* ∈ *E*(*T*)}. The *Robinson-Foulds distance* [32] *(also called the bipartition distance) between trees T* and *T ′* with the same leaf set is RF(*T, T ′*) := |*C*(*T*)*\C*(*T ′*)| + |*C*(*T ′*)*\C*(*T*)|. We extend the definition of RF distance to allow for *T* and *T ′* to have different leaf sets as follows: *RF* (*T, T ′*) := *RF* (*T* |_*X*_, *T ′*|_*X*_), where *X* is the shared leaf set and *t*|_*X*_ denotes the homeomorphic subtree of *t* induced by *X*. Letting *𝒯*^*S*^ denote the set of all phylogenetic trees such that *L*(*T*) = *S* and 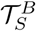 denote the binary trees in *𝒯*_*S*_, then a Robinson-Foulds supertree [2] of a profile *𝒜* is a binary tree

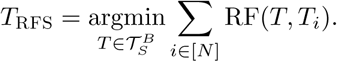

We let RF(*T, 𝒜*) := ∑_*i*∈[*N*]_ RF(*T, T*_*i*_) denote the *RFS score* of *T* with respect to profile *𝒜*. Thus, the **Robinson-Foulds Supertree problem** takes as input the profile *𝒜* and seeks a Robinson-Foulds (RF) supertree for *𝒜*, which we denote by RFS(*𝒜*).

### Split Fit Supertree

The Split Fit (SF) Supertree problem was introduced in [49], and is based on optimizing the number of shared splits (i.e., bipartitions) between the supertree and the source trees. For two trees *T, T ′* with the same leaf set, the *split support* is the number of shared bipartitions, i.e., SF(*T, T ′*) := |*C*(*T*) *∩ C*(*T ′*)|. For trees with different leaf sets, we restrict them to the shared leaf set before calculating the split support. The Split Fit supertree for a profile *𝒜* of source trees, denoted SFS(*𝒜*), is a tree 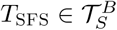 such that

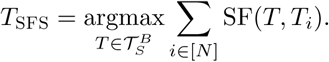

Thus, the split support score of *T* with respect to *𝒜* is SF(*T, 𝒜*) := ∑_*i*∈[*N*]_ SF(*T, T*_*i*_). The **Split Fit Supertree (SFS) problem** takes as input the profile *𝒜* and seeks a Split Fit supertree (the supertree with the maximum split support score), which we denote by SFS(*𝒜*).

*Nomenclature for variants of* RFS *and* SFS *problems*

– The relaxed versions of the problems where we do not require the output to be binary (i.e., we allow *T* ∈ *𝒯*_*S*_) are named Relax-RFS and Relax-SFS.
– We append “-*N*” to the name to indicate that we assume there are *N* source trees. If no number is specified then the number of source trees is unconstrained.
– We append “-B” to the name to indicate that the source trees are required to be binary; hence, we indicate that the source trees are allowed to be non-binary by not appending -B.

Thus, the RFS problem with two binary input trees is RFS-2-B and the relaxed SFS problem with three (not necessarily binary) input trees is Relax-SFS-3.

### Other notation

For any *v* ∈ *V* (*T*), we let *N*_*T*_ (*v*) denote the set of neighbors of *v* in *T*. A tree *T ′* is a *refinement* of *T* iff *T* can be obtained from *T ′* by contracting a set of edges. Two bipartitions *π*_1_ and *π*_2_ of the same leaf set are said to be *compatible* if and only if there exists a tree *T* such that *π*_*i*_ ∈ *C*(*T*), *i* = 1, 2. A bipartition *π* = [*A*|*B*] restricted to a subset *R* is *π*|_*R*_ = [*A ∩ R*| *B ∩ R*]. For a graph *G* and a set *F* of vertices or edges, we use *G* + *F* to represent the graph obtained from adding the set *F* of vertices or edges to *G*, and *G − F* is defined for deletions, similarly.

## 3 Theoretical Results

In this section we establish the main theoretical results, with detailed proofs provided in [50] or in the appendix of the full version on bioRxiv.

### 3.1 Solving RFS and SFS on two trees

#### Lemma 1

*Given an input set 𝒜 of source trees, a tree* 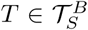 *is an optimal solution for* RFS*(𝒜) if and only if it is an optimal solution for* SFS*(𝒜*).

The main result of this paper is Theorem 1 (correctness is proved later within the main body of the paper, and the running time is established in the Appendix):

#### Theorem 1

*Let 𝒜* = {*T*_1_, *T*_2_*} with S*_*i*_ *the leaf set of T*_*i*_ *(i* = 1, 2*) and X* := *S*_1_ *∩ S*_2_. *The problems* RFS-2*-*B*(𝒜) and* SFS-2*-*B*(𝒜) can be solved in O*(*n*^2^|*X*|) *time, where n* := max{|*S*_1_|, |*S*_2_|}.

#### Exact-RFS-2: Polynomial time algorithm for RFS-2-B and SFS-2-B

The input to Exact-RFS-2 is a pair of binary trees *T*_1_ and *T*_2_. Let *X* denote the set of shared leaves. At a high level, Exact-RFS-2 constructs a tree *T*_init_ that has a central node that is adjacent to every leaf in *X* and to the root of every “rooted extra subtree” (a term we define below) so that *T*_init_ contains all the leaves in *S*. It then modifies *T*_init_ by repeatedly refining it to add specific desired bipartitions, to produce an optimal Split Fit (and optimal Robinson-Foulds) supertree (Figure 3). The bipartitions that are added are defined by a maximum independent set in a bipartite “weighted incompatibility graph” we compute.

##### Additional notation

Let *Π* = 2^*X*^ denote the set of all bipartitions of *X*; any bipartition that splits a single leaf from the remaining |*X*|*−*1 leaves will be called “trivial” and the others will be called “non-trivial”. Let *C*(*T*_1_, *T*_2_, *X*) denote *C*(*T*_1_|_*X*_) *∪ C*(*T*_2_|_*X*_), and let Triv and NonTriv denote the sets of trivial and non-trivial bipartitions in *C*(*T*_1_, *T*_2_, *X*), respectively. We refer to *T*_*i*_|_*X*_, *i* = 1, 2 as **backbone trees** (Figure 2). Recall that we suppress degree-two vertices when restricting a tree *T*_*i*_ to a subset *X* of the leaves; hence, every edge *e* in *T*_*i*_|_*X*_ will correspond to an edge or a path in *T* (see Fig. 1 for an example). We will let *P* (*e*) denote the path associated to edge *e*, and let *w*(*e*) := |*P* (*e*)| (the number of edges in *P* (*e*)). Finally, for *π* ∈ *C*(*T*_*i*_|_*X*_), we define *e*_*i*_(*π*) to be the edge that induces *π* in *T*_*i*_|_*X*_ (Fig. 1).

**Fig. 1:**
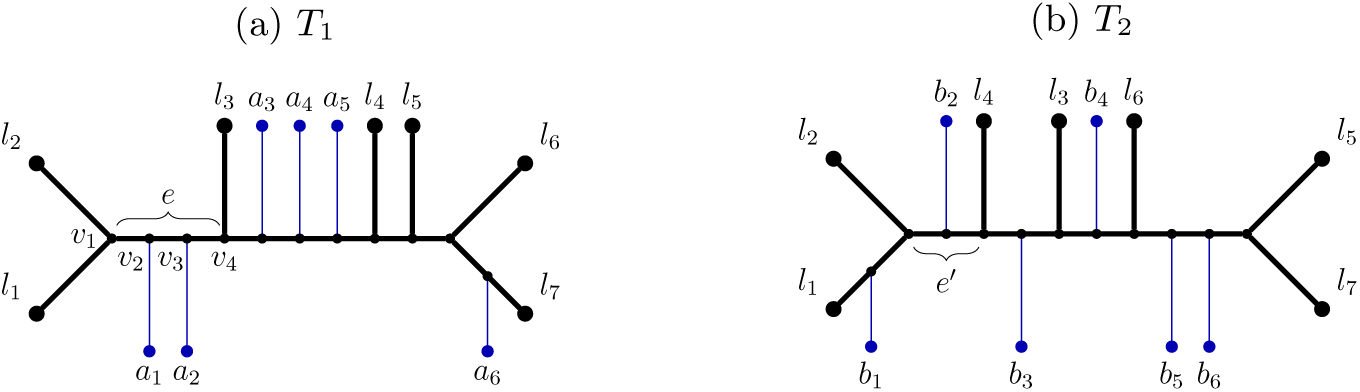
*T*_1_ and *T*_2_ depicted in (a) and (b) have an overlapping leaf set *X* = {*l*_1_, *l*_2_, *…, l*_7_}. Each of *a*_1_, *…, a*_6_ and *b*_1_, *…, b*_6_ can represent a multi-leaf extra subtree. Using indices to represent the shared leaves, let *π* = [12|34567]; then *e*_1_(*π*) = *e* and *e*_2_(*π*) = *e′. P* (*e*) is the path from *v*_1_ to *v*_4_, so *w*(*e*) = 3. 𝒯 ℛ(*e*) = {*a*_1_, *a*_2_}, 𝒯 ℛ(*e′*) = {*b*_2_}, so *𝒯 ℛ\**(*π*) = {*a*_1_, *a*_2_, *b*_2_}. Let *A* = {1, 2, 3*}, B* = {4, 5, 6, 7}. Ignoring the trivial bipartitions, we have ℬ𝒫(*A*) = {[12|34567]} and ℬ𝒫(*B*) = {[1234|567], [12345|67], [12346|57]}. 𝒯 ℛ𝒮(*A*) = {*a*_1_, *a*_2_, *b*_2_} and 𝒯 ℛ𝒮(*B*) = {*b*_4_, *b*_5_, *b*_6_}.

**Fig. 2:**
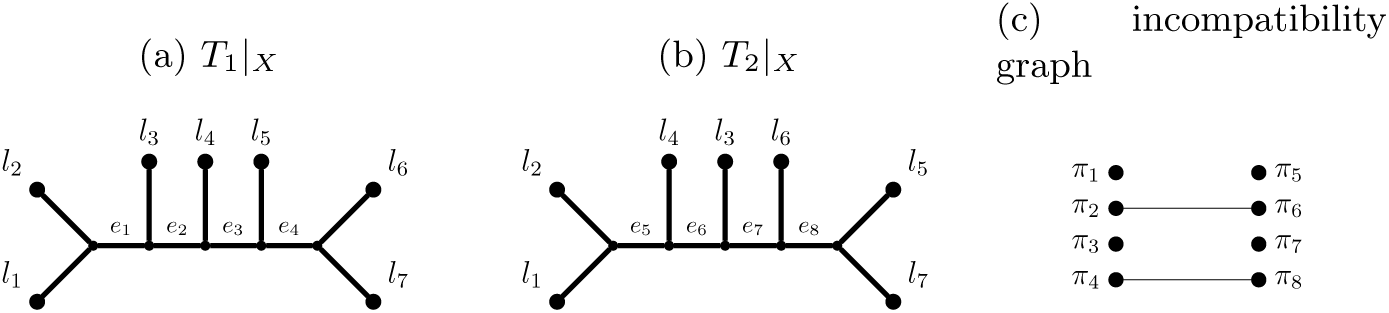
We show (a) *T*_1_|_*X*_, (b) *T*_2_|_*X*_, and (c) their incompatibility graph, based on the trees *T*_1_ and *T*_2_ in Figure 1 (without the trivial bipartitions). Each *π*_*i*_ is the bipartition induced by *e*_*i*_, and the weights for *π*_1_, *…, π*_8_ are 3, 4, 1, 1, 2, 2, 2, 3, in that order. We note that *π*_1_ and *π*_5_ are the same bipartition, but they have different weights as they are induced by different edges; similarly for *π*_3_ and *π*_7_. The maximum weight independent set in this graph has all the isolated vertices (*π*_1_, *π*_3_, *π*_5_, *π*_7_) and also *π*_2_, *π*_8_, and so has total weight 15.

The next concept we introduce is the set of **extra subtrees**, which are rooted subtrees of *T*_1_ and *T*_2_, formed by deleting *X* and all the edges and vertices on paths between vertices in *X* (i.e., we delete *T*_*i*_|_*X*_ from *T*_*i*_). Each component in *T*_*i*_ *− T*_*i*_|_*X*_ is called an **extra subtree** of *T*_*i*_, and note that the extra subtree *t* is naturally seen as rooted at the unique vertex *r*(*t*) that is adjacent to a vertex in *T*_*i*_|_*X*_. Thus, Extra(*T*_*i*_) = {*t* | *t* is a component in *T*_*i*_ *− T*_*i*_|_*X*_}.

We can now define the initial tree *T*_init_ computed by Exact-RFS-2: *T*_init_ has a center node that is adjacent to every *x* ∈ *X* and also to the root *r*(*t*) for every extra subtree *t* ∈ Extra(*T*_1_) ∪ Extra(*T*_2_). Note that *T*_init_ has a leaf for every element in *S*, and that *T*_init_|_*S*_*i* is a contraction of *T*_*i*_, formed by collapsing all the edges in the backbone tree *T*_*i*_|_*X*_.

We say that an extra subtree *t* is **attached to edge** *e* ∈ *E*(*T*_*i*_|_*X*_) if the root of *t* is adjacent to an internal node of *P* (*e*), and we let 𝒯 ℛ(*e*) denote the set of such extra subtrees attached to edge *e*. Similarly, if *π* ∈ *C*(*T*_1_, *T*_2_, *X*), we let 𝒯 ℛ^*\**^(*π*) refer to the set of extra subtrees that attach to edges in a backbone tree that induce *π* in either *T*_1_|_*X*_ or *T*_2_|_*X*_. For exuample, if both trees *T*_1_ and *T*_2_ contribute extra subtrees to *π*, then 𝒯 ℛ^*\**^(*π*) :=∪ _*i*∈[2]_ 𝒯 ℛ(*e*_*i*_(*π*)).

For any *Q* ⊆ *X*, we let ℬ𝒫_*i*_(*Q*) denote the set of bipartitions in *C*(*T*_*i*_|_*X*_) that have one side being a strict subset of *Q*, and we let 𝒯 ℛ 𝒮_*i*_(*Q*) denote the set of extra subtrees associated with these bipartitions. In other words, ℬ𝒫_*i*_(*Q*) := {[*A*|*B*] ∈ *C*(*T*_*i*_|_*X*_) | *A* ⊊ *Q* or *B* ⊊ *Q*}, and 𝒯 ℛ 𝒮_*i*_(*Q*) :=∪_*π∈ℬ𝒫*_*i*_(*Q*)_ 𝒯 ℛ(*e*_*i*_(*π*)). Intuitively, 𝒯 ℛ 𝒮_*i*_(*Q*) denotes the set of extra subtrees in *T*_*i*_ that are “on the side of *Q*”. By Corollary 2, which appears in the Appendix, for any *π* = [*A*|*B*] ∈ *C*(*T*_*i*_|_*X*_), ℬ𝒫_*i*_(*A*)*∪ℬ𝒫*_*i*_(*B*) is the set of bipartitions in *C*(*T*_*i*_|_*X*_) that are compatible with *π*. Finally, let ℬ𝒫(*Q*) = ℬ𝒫_1_(*Q*)*∪ℬ𝒫*_2_(*Q*), and 𝒯 ℛ 𝒮(*Q*) = 𝒯 ℛ 𝒮_1_(*Q*)*∪ 𝒯 RS*_2_(*Q*). We give an example for these terms in Figure 1.

The *incompatibility graph* of a set of trees, each on the same set of leaves, has one vertex for each bipartition in any tree (and note that bipartitions can appear more than once) and edges between bipartitions if they are incompatible (see [30]). We compute a **weighted incompatibility graph** for the pair of trees *T*_1_|_*X*_ and *T*_2_|_*X*_, in which the weight of the vertex corresponding to bipartition *π* appearing in tree *T*_*i*_|_*X*_ is *w*(*e*_*i*_(*π*)), as defined previously. Thus, if a bipartition is common to the two trees, it produces two vertices in the weighted incompatibility graph, and each vertex has its own weight (Figure 2).

We divide 𝒞 = *C*(*T*_1_) *∪ C*(*T*_2_) into two sets: *Π*_*X*_ = {[*A*|*B*] ∈ *C* | *A ∩ X ≠ ∅* and *B ∩ X ≠ ∅*}, and *Π*_*Y*_ = {[*A*|*B*] ∈ *C* | *A ∩ X* = *∅* or *B ∩ X* = *∅*}. Intuitively, *Π*_*X*_ is the set of bipartitions from the input trees that are induced by edges in the minimal subtree of *T*_1_ or *T*_2_ spanning *X*, and *Π*_*Y*_ are all the other input tree bipartitions. We define *p*_*X*_ (·) and *p*_*Y*_ (·) on trees *T* ∈ *𝒯*_*S*_ by:

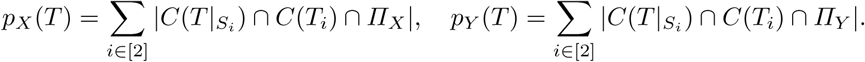

###### Algorithm 1 Exact-RFS-2: Computing a Robinson-Foulds supertree of two trees (see Figure 3)

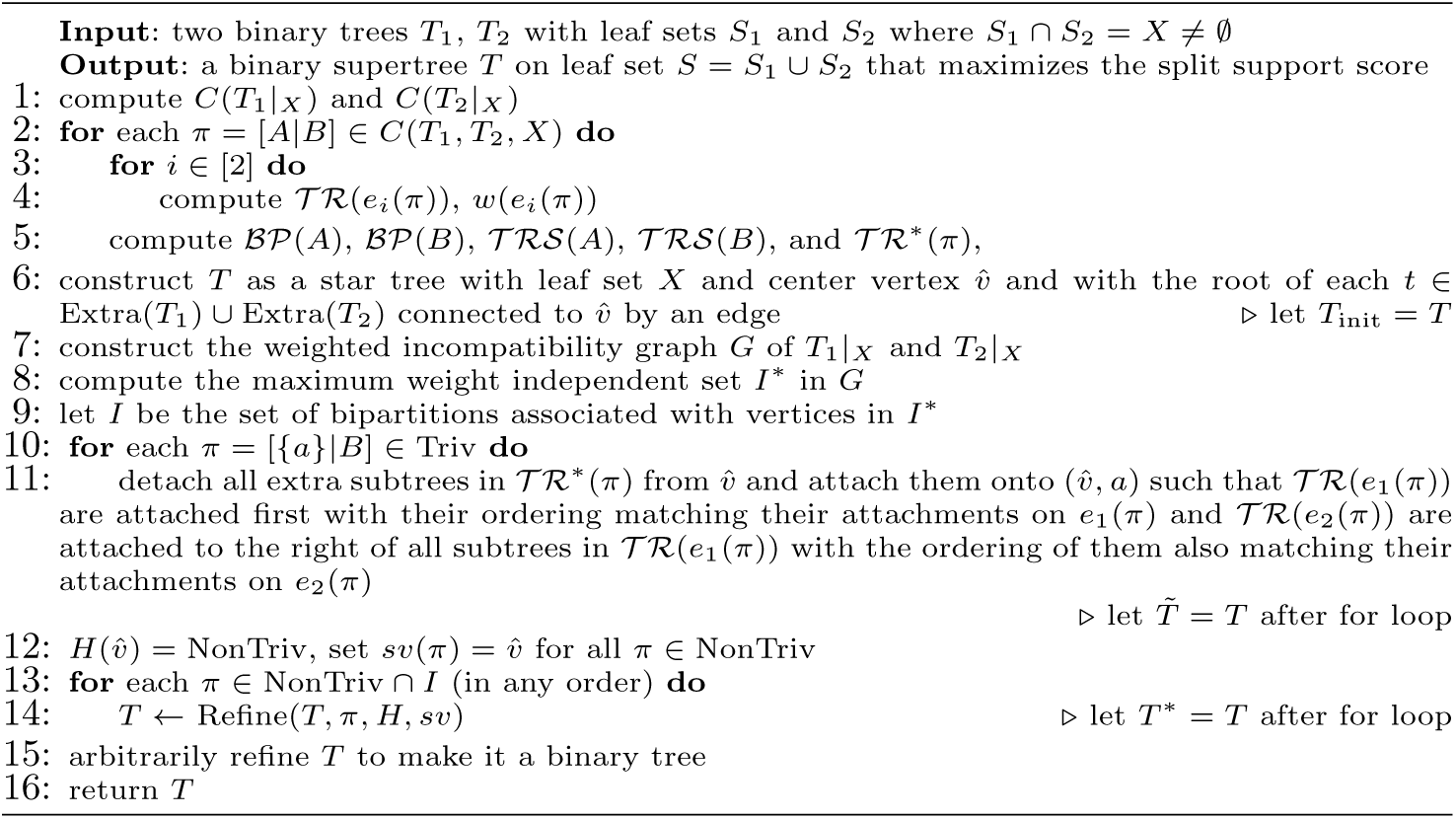

**Fig. 3:**
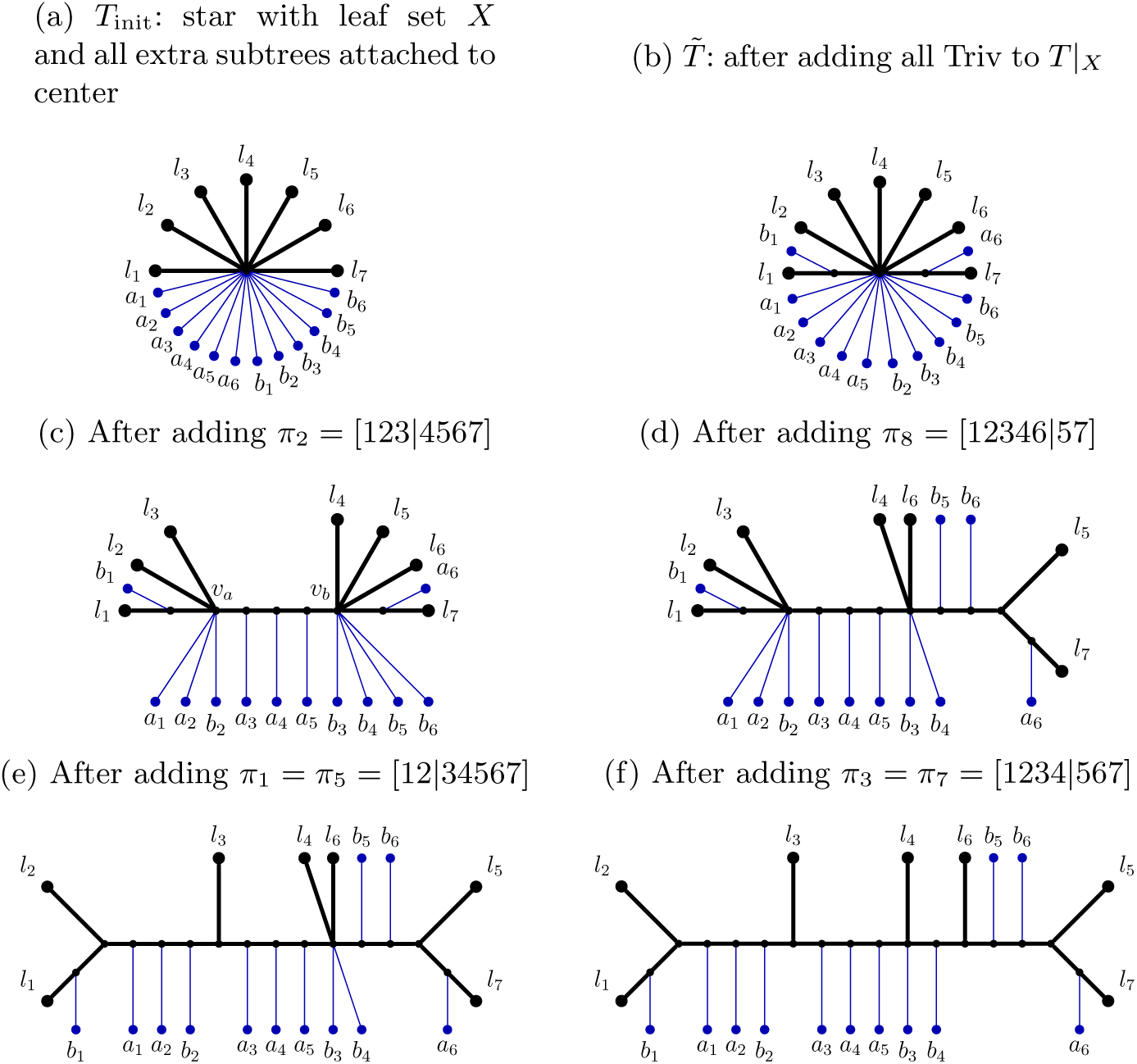
Algorithm 1 working on *T*_1_ and *T*_2_ from Figure 1 as source trees; the indices of leaves in *X* = {*l*_1_, *l*_2_, *…, l*_7_} represent the leaves and the notation of *π*_1_, *…, π*_8_ is from Figure 2. In (a) to (f), the *p*_*X*_ (·) score of the trees are 14, 16, 20, 23, 27, 29, in that order. We explain how the algorithm obtain the tree in (c) from 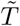 by adding *π*_2_ = [123|4567] to the backbone of 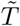. Let *A* = {*l*_1_, *l*_2_, *l*_3_} and *B* = {*l*_4_, *l*_5_, *l*_6_, *l*_7_}. The center vertex *c* of 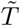 is split into two vertices *v*_*a*_, *v*_*b*_ with an edge between them. Then all neighbors of *c* between *c* and *A* are made adjacent to *v*_*a*_ while the neighbors between *c* and *B* are made adjacent to *v*_*b*_. All neighbors of *c* which are roots of extra subtrees are moved around such that all extra subtrees in 𝒯 ℛ*(*π*_2_) are attached onto (*v*_*a*_, *v*_*b*_); all extra subtrees in 𝒯 ℛ 𝒮(*A*) = {*a*_1_, *a*_2_, *b*_2_} are attached to *v*_*a*_ and all extra subtrees in 𝒯 ℛ 𝒮(*B*) = {*b*_4_, *b*_5_, *b*_6_} are attached to *v*_*b*_. We note that in this step, *b*_3_ can attach to either *v*_*a*_ or *v*_*b*_ because it is not in 𝒯 ℛ 𝒮(*A*) or 𝒯 ℛ 𝒮(*B*). However, when obtaining the tree in (d) from the tree in (c), *b*_3_ can only attach to the left side because for *A′* = {*l*_1_, *l*_2_, *l*_3_, *l*_4_, *l*_6_}, [124|3567] ∈ ℬ𝒫(*A′*) and thus *b*_3_ ∈ 𝒯 ℛ 𝒮(*A′*).

Note that *p*_*X*_ (*T*) and *p*_*Y*_ (*T*) decompose the split support score of *T* into the score contributed by bipartitions in *Π*_*X*_ and the score contributed by bipartitions in *Π*_*Y*_ ; thus, the split support score of *T* with respect to *T*_1_, *T*_2_ is *p*_*X*_ (*T*) + *p*_*Y*_ (*T*). As we will show, the two scores can be maximized independently and we can use this observation to refine *T*_init_ so that it achieves the optimal total score.

##### Overview of Exact-RFS-2

Exact-RFS-2 (Algorithm 1) has four phases. In the pre-processing phase (lines 1–5), it computes the weight function *w* and the mappings 𝒯 ℛ, 𝒯 ℛ^*\**^, ℬ𝒫, and 𝒯 ℛ 𝒮 for use in latter parts of Algorithm 1 and Algorithm 2. In the initial construction phase (line 6), it constructs a tree *T*_init_ (as described earlier), and we note that *T*_init_ maximizes *p*_*Y*_ (·) score (Lemma 2). In the refinement phase (lines 7–14), it refines *T*_init_ so that it attains the maximum *p*_*X*_ (·) score. In the last phase (line 15), it arbitrarily refines *T* to make it binary. The refinement phase begins with the construction of a weighted incompatibility graph *G* of *T*_1_|_*X*_ and *T*_2_|_*X*_ (see Figure 2). It then finds a maximum weight independent set of *G* that defines a set *I ⊆ C*(*T*_1_, *T*_2_, *X*) of compatible bipartitions of *X*. Finally, it uses these bipartitions of *X* in *I* to refine *T*_init_ to achieve the optimal *p*_*X*_ (·) score, by repeatedly applying Algorithm 2 for each *π* ∈ *I* (and we note that the order does not matter). See Figure 3 for an example of Exact-RFS-2 given two input source trees.

Algorithm 2 refines the given tree *T* on leaf set *S* with bipartitions on *X* from *C*(*T*_1_, *T*_2_, *X*) *\ C*(*T* |_*X*_). Given bipartition *π* = [*A*|*B*] on *X*, Algorithm 2 produces a refinement *T ′* of *T* such that *C*(*T ′*|_*S*_*i*) = *C*(*T* |_*S*_*i*) *∪ {π′* ∈ *C*(*T*_*i*_) | *π′*|_*X*_ = *π*} for both *i* = 1, 2. To do this, we first find the unique vertex *v* such that no component of *T − v* has leaves from both *A* and *B*. We create two new vertices *v*_*a*_ and *v*_*b*_ with an edge between them. We divide the neighbor set of *v* into three sets: *N*_*A*_ is the set of neighbors that split *v* from leaves in *A, N*_*B*_ is the set of neighbors that split *v* from leaves in *B*, and *N*_other_ contains the remaining neighbors. Then, we make vertices in *N*_*A*_ adjacent to *v*_*a*_ and vertices in *N*_*B*_ adjacent to *v*_*b*_. We note that *N*_other_ = *∅* if *X* = *S* and thus there are no extra subtrees. In the case where *X* ≠ *S, N*_other_ contains the roots of the extra subtrees adjacent to *v* and we handle them in four different cases to make *T ′* include the desired bipartitions:

###### Algorithm 2 Refine

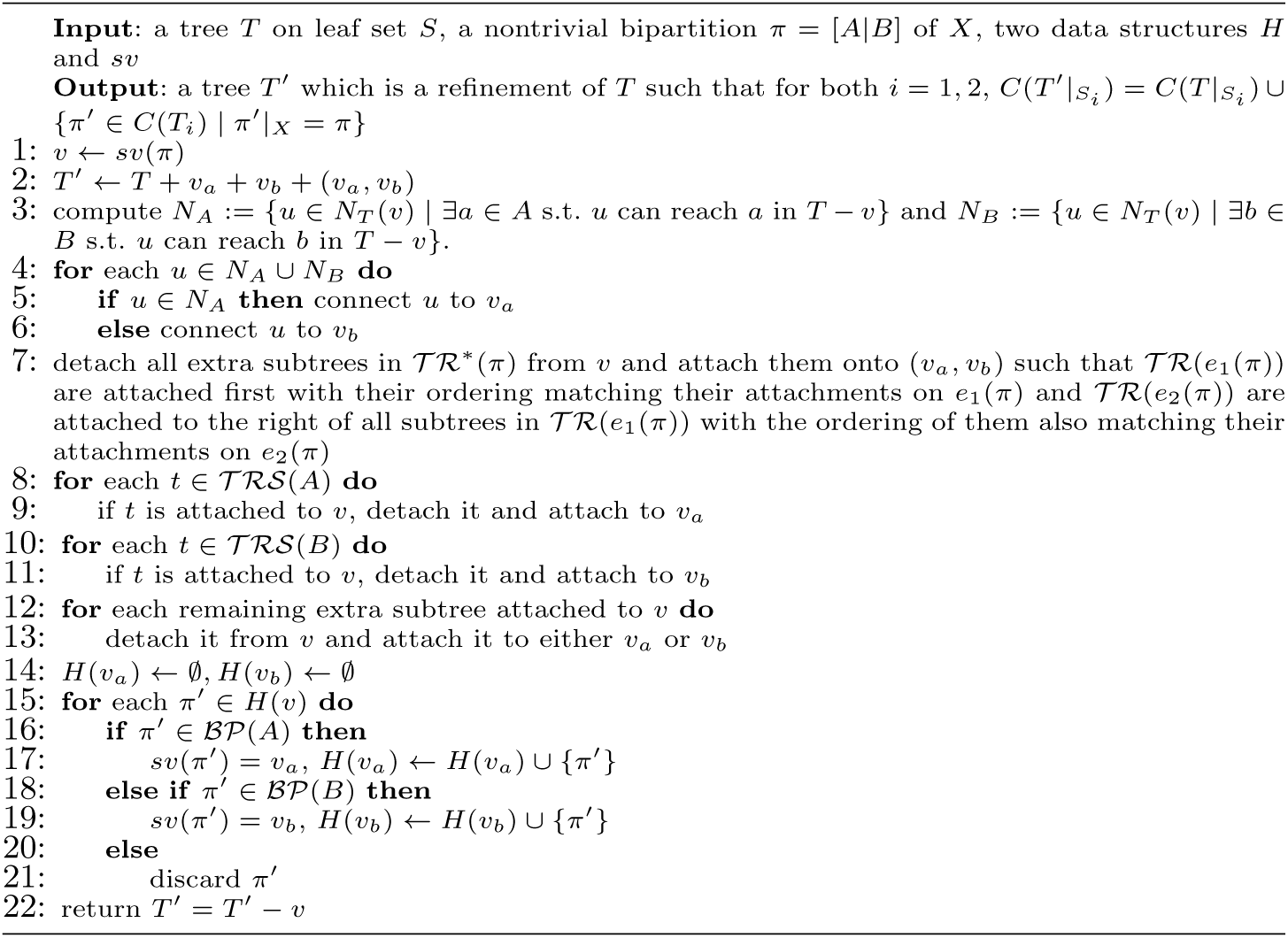

– those vertices that root extra subtrees in 𝒯 ℛ^*\**^(*π*) are moved onto the edge (*v*_*a*_, *v*_*b*_) (by subdividing the edge to create new vertices, and then making these vertices adjacent to the new vertices)
– those vertices that root extra subtrees in 𝒯 ℛ 𝒮(*A*) are made adjacent to *v*_*a*_
– those that root extra subtrees in 𝒯 ℛ 𝒮(*B*) are made adjacent to *v*_*b*_
– the remaining vertices can be made adjacent to either *v*_*a*_ or *v*_*b*_

Algorithms 1 and 2 also use two data structures (functions) *H* and *sv*: (1) For a given node *v* ∈ *V* (*T*), *H*(*v*) ⊆ *C*(*T*_1_, *T*_2_, *X*) is the set of bipartitions of *X* that can be added to *T* |_*X*_ by refining *T* |_*X*_ at *v*, and (2) Given *π* ∈ *C*(*T*_1_, *T*_2_, *X*), *sv*(*π*) = *v* means ∃*T ′*, a refinement of *T* at *v*, so that *C*(*T ′*|_*X*_) = *C*(*T* |_*X*_) *∪ {π*}.

###### Lemma 2

*For any tree T* ∈ *𝒯*_*S*_, *p*_*Y*_ (*T*) *≤* |*Π*_*Y*_ |. *In particular, let T*_init_ *be the tree defined in line 6 of Algorithm 1. Then, p*_*Y*_ (*T*_init_) = |*Π*_*Y*_ |.

Lemma 2 formally states that the tree *T*_init_ we build in line 6 of Exact-RFS-2 (Algorithm 1) maximizes the *p*_*Y*_ (·) score. This lemma is true because there are only |*Π*_*Y*_ | bipartitions that can contribute to *p*_*Y*_ (·) and *T*_init_ contains all of them by construction. We define the function *w*^*\**^ : *Π →* ℕ_*≥*0_ as follows:

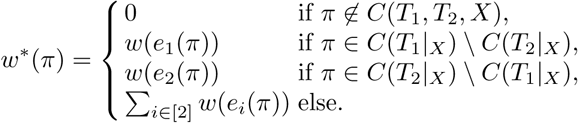

For any set *F* of bipartitions, we let *w*^*\**^(*F*) =∑ _*π∈F*_ *w*^*\**^(*π*).

###### Lemma 3

*Let π* = [*A*|*B*] ∈ *Π. Let T* ∈ *𝒯*_*S*_ *be any tree with leaf set S such that π ∉ C*(*T* |_*X*_) *but π is compatible with C*(*T* |_*X*_). *Let T ′ be a refinement of T such that for all π′* ∈ *C*(*T ′*|_*S*_*i*)*\C*(*T* |_*S*_*i*) *for some i* ∈ [2], *π′*|_*X*_ = *π. Then, p*_*X*_ (*T ′*) *− p*_*X*_ (*T*) *≤ w*^*\**^(*π*).

###### Lemma 4

*For any compatible set F ⊆ Π, let T* ∈ *𝒯*_*S*_ *be any tree with leaf set S such that C*(*T* |_*X*_) = *F*. *Then p*_*X*_ (*T*) *≤ w*^*\**^(*F*).

Lemma 3 shows that *w*^*\**^(*π*) represents the maximum potential increase in *p*_*X*_ (·) as a result of adding bipartition *π* to *T*|_*X*_. The proof of Lemma 3 follows the idea that for any bipartition *π* of *X*, there are at most *w*^*\**^(*π*) edges in either *T*_1_ or *T*_2_ whose induced bipartitions become *π* when restricted to *X*. Therefore, by only adding *π* to *T* |_*X*_, at most *w*^*\**^(*π*) more bipartitions get included in 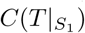 or 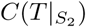 so that they contribute to the increase of *p*_*X*_ (*T*). The proof of Lemma 4 uses Lemma 3 repeatedly by adding the compatible bipartitions to the tree in an arbitrary order.

###### Proposition 1

*Let* 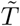 *be the tree constructed after line 11 of Algorithm 1, then* 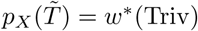.

The proof naturally follows by construction (Line 8 of Algorithm 1), and implies that the algorithm adds the trivial bipartitions of *X* (which are all in *I*) to *T* |_*X*_ so that *p*_*X*_ (*T*) reaches the full potential of adding those trivial bipartitions.

###### Lemma 5

*Let T be a supertree computed within Algorithm 1 at line* 14 *immediately before a refinement step. Let π* = [*A*|*B*] ∈ NonTriv *∩ I. Let T ′ be a refinement of T obtained from running Algorithm 2 with supertree T, bipartition π, and the auxiliary data structures H and sv. Then, p*_*X*_ (*T ′*) *− p*_*X*_ (*T*) = *w*^*\**^(*π*).

The idea for the proof of Lemma 5 is that for any non-trivial bipartition *π* ∈ *I* of *X* to be added to *T* |_*X*_, Algorithm 2 is able to split the vertex correctly and move extra subtrees around in a way such that each bipartition in *T*_1_ or *T*_2_ that is induced by an edge in *P* (*e*_1_(*π*)) or *P* (*e*_2_(*π*)), which is not in 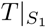 or 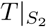 before the refinement, becomes present in 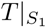 or 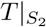 after the refinement. Since there are exactly *w*^*\**^(*π*) such bipartitions, they increase *p*_*X*_ (·) by *w*^*\**^(*π*).

###### Proposition 2

*Let G be the weighted incompatibility graph on T*_1_|_*X*_ *and T*_2_|_*X*_, *and let I be the set of bipartitions associated with vertices in I*^*\**^, *which is a maximum weight independent set of G. Let F be any compatible subset of C*(*T*_1_, *T*_2_, *X*). *Then w*^*\**^(*I*) *≥ w*^*\**^(*F*).

We now restate and prove Theorem 1:

###### Theorem 1

*Let 𝒜* = {*T*_1_, *T*_2_} *with S*_*i*_ *the leaf set of T*_*i*_ *(i* = 1, 2*) and X* := *S*_1_ *∩ S*_2_. *The problems* RFS-2*-*B*(A) and* SFS-2*-*B*(A) can be solved in O*(*n*^2^|*X*|) *time, where n* := max{|*S*_1_|, |*S*_2_|}.

*Proof*. First we claim that *p*_*X*_ (*T* ^*\**^) *≥ p*_*X*_ (*T*) for any tree *T* ∈ *T*_*S*_, where *T* ^*\**^ is defined as from line 14 of Algorithm 1. Fix arbitrary *T* ∈ *𝒯*_*S*_ and let *F* = *C*(*T* |_*X*_). Then by Lemma 4, *p*_*X*_ (*T*) *≤ w*^*\**^(*F*). We know that *w*^*\**^(*π*) = 0 for any *π ∉ C*(*T*_1_, *T*_2_, *X*), so *w*^*\**^(*F*) = *w*^*\**^(*F ∩ C*(*T*_1_, *T*_2_, *X*)) and thus *p*_*X*_ (*T*) *≤ w*^*\**^(*F ∩ C*(*T*_1_, *T*_2_, *X*)). Since *F ∩ C*(*T*_1_, *T*_2_, *X*) is a compatible subset of *C*(*T*_1_, *T*_2_, *X*), we have *w*^*\**^(*F ∩ C*(*T*_1_, *T*_2_, *X*)) *≤ w*^*\**^(*I*) by Proposition 2. Then *p*_*X*_ (*T*) *≤ w*^*\**^(*I*). Since Triv ⊆ *C*(*T*_1_|_*X*_) *∩ C*(*T*_2_|_*X*_) ⊆ *I*, we have *I* = (NonTriv *∩ I*) ∪ (Triv *∩ I*) = (NonTriv *∩ I*) ∪ Triv. Therefore, by Proposition 1 and Lemma 5, we have

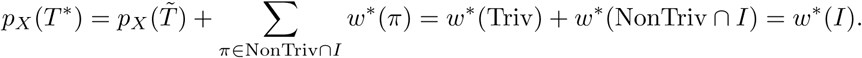

Therefore, *p*_*X*_ (*T* ^*\**^) = *w*^*\**^(*I*) *≥ p*_*X*_ (*T*).

From Lemma 2 and the fact that a refinement of a tree never decreases *p*_*X*_ (·) and *p*_*Y*_ (·), we also know that *p*_*Y*_ (*T* ^*\**^)*≥ p*_*Y*_ (*T*_init_) *≥ p*_*Y*_ (*T*) for any tree *T* ∈ *𝒯*_*S*_. Since for any *T* ∈ *𝒯*_*S*_, SF(*T, 𝒜*) = *p*_*X*_ (*T*) + *p*_*Y*_ (*T*), *T* ^*\**^ achieves the maximum split support score with respect to *A* among all trees in *𝒯*_*S*_. Thus, *T* ^*\**^ is a solution to Relax–SFS-2-B (Corollary 1). If *T* ^*\**^ is not binary, Algorithm 1 arbitrarily resolves every high degree node in *T* ^*\**^ until it is a binary tree and then returns a tree that achieves the maximum split support score among all binary trees of leaf set *S*. See the Appendix for the running time analysis. □

###### Corollary 1

*Let A* = {*T*_1_, *T*_2_} *with S*_*i*_ *the leaf set of T*_*i*_ *(i* = 1, 2*) and X* := *S*_1_*∩ S*_2_. Relax*–*SFS-2*-*B *can be solved in O*(*n*^2^|*X*|) *time, where n* := max{|*S*_1_|, |*S*_2_|}.

###### Lemma 6

RFS-3, SFS-3, *and* Relax*–*SFS-3 *are all* **NP***-hard*.

### 3.2 DACTAL-Exact-RFS-2

Let *F* be a model of evolution (e.g., GTR) for which statistically consistent methods exist, and we have some data (e.g., sequences) generated by the model and wish to construct a tree. We construct an initial estimate of the tree, divide the dataset into two overlapping subsets (by removing an edge *e*, letting *X* be the set of the *p* nearest leaves to *e*, and letting the subsets be *A* ∪ *X* and *B* ∪ *X*, where *π*_*e*_ = [*A*|*B*]), re-estimate trees on the subsets (perhaps using a recursive approach that is statistically consistent), and then combine the trees together using Exact-RFS-2. We call this the DACTAL-Exact-RFS-2 pipeline, due to its similarity to the DACTAL pipeline [26].

Before we prove that DACTAL-Exact-RFS-2 can enable statistically consistent pipelines, we begin with some definitions. Given a tree *T* and an internal edge *e* in *T*, the deletion of the edge *e* and its endpoints defines four subtrees. A **short quartet around** *e* is a set of four leaves, one from each subtree, selected to be closest to the edge. Note that due to ties, there can be multiple short quartets around some edges. The set of short quartets for a tree *T* is the set of all short quartets around the edges of *T*. The **short quartet trees of** *T* is the set of quartet trees on short quartets induced by *T*. It is well known that the short quartet trees of a tree *T* define *T*, and furthermore *T* can be computed from this set in polynomial time [11–13].

#### Lemma 7

*Let T be a binary tree on leaf set S and let A, B ⊆ S. Let T*_*A*_ = *T* |_*A*_ *and T*_*B*_ = *T* |_*B*_ *(i.e*., *T*_*A*_ *and T*_*B*_ *are correct induced subtrees). If every short quartet tree is induced in T*_*A*_ *or in T*_*B*_, *then T is the unique compatibility supertree for T*_*A*_ *and T*_*B*_ *and Exact-2-RFS(T*_*A*_, *T*_*B*_) = *T*.

*Proof*. Because *T*_*A*_ and *T*_*B*_ are true trees, it follows that *T* is a compatibility supertree for *T*_*A*_ and *T*_*B*_. Furthermore, because every short quartet tree appears in at least one of these trees, *T* is the unique compatibility supertree for *T*_*A*_ and *T*_*B*_ (by results from [12, 13], mentioned above). Finally, because *T* is a compatibility supertree, the RFS score of *T* with respect to *T*_*A*_, *T*_*B*_ is 0, which is the best possible. Since Exact-2-RFS solves the RFS problem on two binary trees, Exact-2-RFS returns *T* given input *T*_*A*_ and *T*_*B*_. □

Thus, Exact-2-RFS is guaranteed to return the true tree when given two correct trees that have sufficient overlap (in that all short quartets are included). We continue with proving that these pipelines are statistically consistent.

#### Theorem 2

*The DACTAL-Exact-RFS-2 pipeline is statistically consistent under any model F for which statistically consistent tree estimation methods exist*.

*Proof*. Let *Φ* be the sequence evolution model. To establish statistical consistency of the pipeline, we need to prove that as the amount of data increases the tree that is returned by the pipeline converge to the true tree. Hence, let *F* be the method used to compute the starting tree and let *G* be the method used to compute the subset trees. Because *F* is statistically consistent under *Φ*, it follows that as the amount of data increases, the starting tree computed by *F* will converge to the true tree *T*. Now consider the decomposition into two sets produced by the algorithm, when applied to the true tree. Let *e* be the edge that is deleted and let the four subtrees around *e* have leaf sets *A*_1_, *A*_2_, *B*_1_, and *B*_2_. Note in particular that all the leaves appearing in any short quartet around *e* are placed in the set *X*. Now, subset trees are computed using *G* on *A*_1_ *∪ A*_2_ *∪ X* and *B*_1_ *∪ B*_2_ *∪ X*, which we will refer to as *T*_*A*_ and *T*_*B*_, respectively. Since *G* is statistically consistent, as the amount of data increases, *T*_*A*_ converges to the true tree on its leaf set (*T* |_*L*(*TA*)_) and *T*_*B*_ converges to the true tree on its leaf set 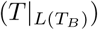. When *T*_*A*_ and *T*_*B*_ are equal to the true trees on their leaf sets, then every short quartet tree of *T* is in *T*_*A*_ or *T*_*B*_, and by Lemma 7, *T* is the only compatibility supertree for *T*_*A*_ and *T*_*B*_ and Exact-2-RFS(*T*_*A*_, *T*_*B*_) returns *T*. □

Hence, DACTAL+Exact-2-RFS is statistically consistent under all standard molecular sequence evolution models and also under the MSC+GTR model [33, 45] where gene trees evolve within species trees under the multi-species coalescent model (which addresses gene tree discordance due to incomplete lineage sorting [19]) and then sequences evolve down each gene tree under the GTR model.

Note that all that is needed for *F* and *G* to guarantee that the pipeline is statistically consistent is that they should be statistically consistent under *Φ*. However, for the sake of improving empirical performance, *F* should be fast so that it can run on the full dataset but *G* can be more freely chosen, since it will only be run on smaller datasets. Indeed, the user can specify the size of the subsets that are analyzed, with smaller datasets enabling the use of more computationally intensive methods.

For example, when estimating trees under the GTR [42] model, *F* could be a fast but statistically consistent distance-based method such as neighbor joining [34] and *G* could be RAxML [37], a leading maximum likelihood method. For the MSC+GTR model, *F* and *G* could be polynomial time summary methods (i.e., methods that estimate the species tree by combining gene trees), with *F* being ASTRID [43] (a very fast summary method) and *G* being ASTRAL [23, 24, 51], which is slower than ASTRID but often more accurate. However, if the subsets are chosen to be very small, then other choices for *G* include StarBeast2 [28], a Bayesian method for co-estimating gene trees and species trees under the MSC+GTR model.

## 4 Experiments and Results

We performed two experiments: Experiment 1, where we used Exact-2-RFS within a divide-and-conquer strategy for large scale phylogenomic species tree estimation where gene trees differ from the species tree due to Incomplete Lineage Sorting (ILS), and Experiment 2, where we used Exact-2-RFS as part of a greedy heuristic, GreedyRFS, for large-scale supertree estimation.

### 4.1 Experiment 1: Phylogenomic species tree estimation

In this experiment, the input is a set of gene trees that can differ from the species tree due to Incomplete Lineage Sorting [19], ASTRAL [23, 24, 51] is used to construct species trees on the two overlapping subsets in the DACTAL pipeline described above, and the two overlapping estimated species trees are then merged together using either Exact-2-RFS or FastRFS. Because the divide-and-conquer strategy produces two source trees, the RFS criterion score for Exact-2-RFS cannot be worse than the score obtained by FastRFS; here we examine the degree of improvement. The simulation protocol produced datasets with high variability (especially for small numbers of genes), so that there was substantial range in the optimal criterion scores for 25 and 100 genes (Figure 4). On average, Exact-2-RFS produces better RFS scores than FastRFS for all numbers of genes, and strictly dominates FastRFS for 1000 genes (Fig. 4), showing that divide-and-conquer pipelines are improved using Exact-2-RFS compared to FastRFS.

**Fig. 4:**
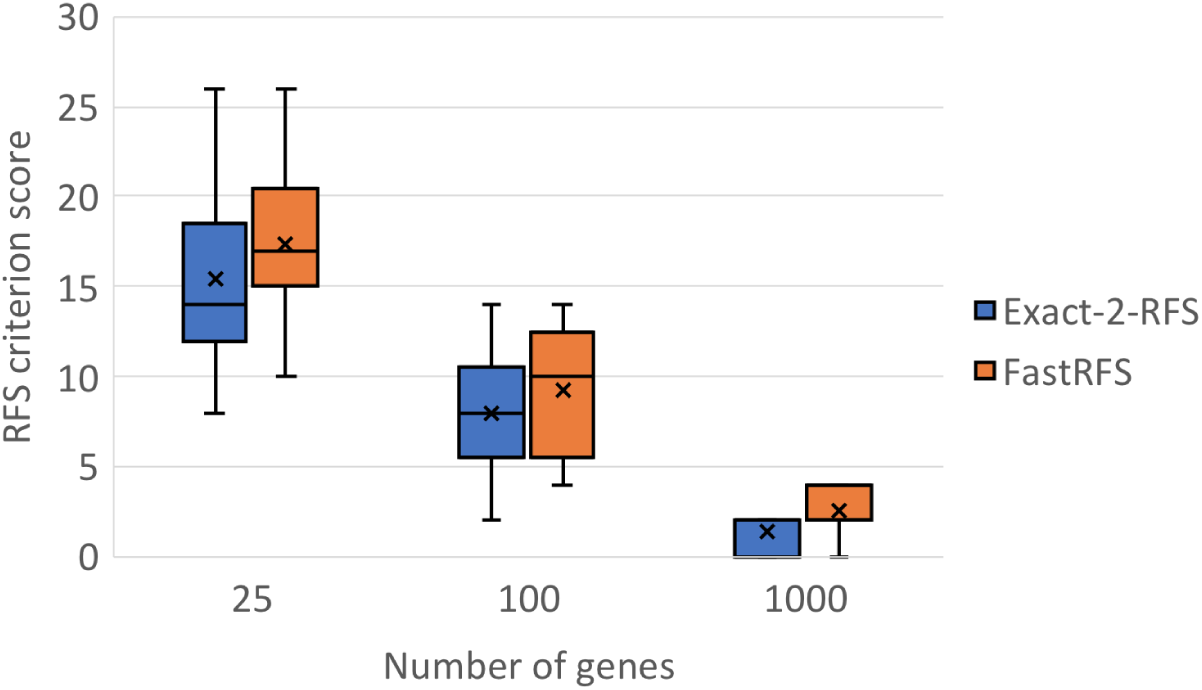
Results for Experiment 1: Exact-2-RFS has better RFS criterion scores than FastRFS (lower is better) in ILS-based species tree estimation (using ASTRAL-III [51], for 501 species with varying numbers of genes.

### 4.2 Experiment 2: Exploring GreedyRFS for supertree estimation

We developed GreedyRFS, a greedy heuristic that takes a profile *A* as input, and then merges pairs of trees until all the trees are merged into a single tree. The choice of which pair to merge follows the technique used in SuperFine [41] for computing the Strict Consensus Merger, which selects the pair that maximizes the number of shared taxa between the two trees (other techniques could be used, potentially with better accuracy [15]). Thus, GreedyRFS is identical to Exact-2-RFS when the profile has only two trees.

We use a subset of the SMIDgen [39] datasets with 500 species and varying numbers of source trees (each estimated using maximum likelihood heuristics) that have been used to evaluate supertree methods in several studies [27, 39–41, 44]. See Appendix (in the full version of the paper on arXiv) for full details of this study.

We explored the impact of changing the number of source trees. The result for two source trees is predicted by theory (i.e., GreedyRFS is the same as Exact-2-RFS for two source trees, and so is guaranteed optimal for this case), but even when the number of source trees was greater than two, GreedyRFS dominated FastRFS in terms of criterion score, provided that the number of source trees was not too large (Fig. 5).

**Fig. 5:**
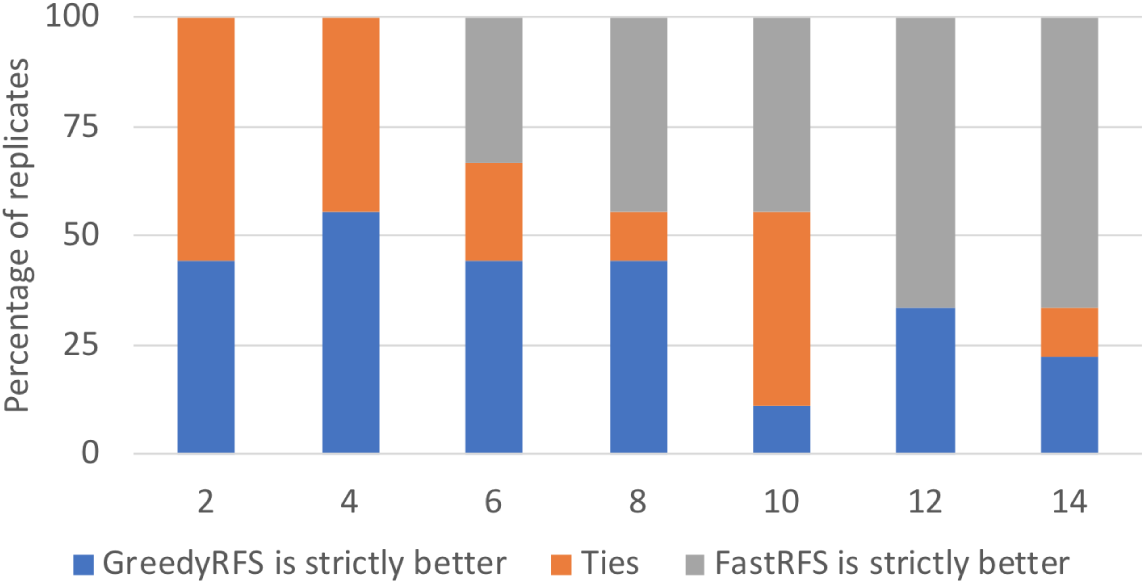
Results for Experiment 2: The percentage of datasets (*y*-axis) that each method (FastRFS and GreedRFS) ties with or is strictly better than the other in terms of RFS criterion score is shown for varying numbers of source trees (*x*-axis), based on nine replicate supertree 500-leaf 20% scaffold datasets (from [41])

This establishes that the advantage in criterion score is not limited to the case of two source trees, suggesting that using Exact-2-RFS within GreedyRFS (or some other heuristics) may be useful for supertree estimation more generally.

## 5 Conclusions

The main contribution of this paper is Exact-2-RFS, a polynomial time algorithm for the Robinson-Foulds Supertree (RFS) of two trees that enables divide-and-conquer pipelines to be provably statistically consistent under sequence evolution models (e.g., GTR [42] and MSC+GTR [33]). Our experimental study showed that Exact-2-RFS dominates the leading RFS heuristic, FastRFS, when used within divide-and-conquer species tree estimation using genome-scale datasets, a problem of increasing importance in biology. We also showed that a greedy heuristic using Exact-2-RFS produced better criterion scores than FastRFS when the number of source trees was small to moderate, showing the potential for Exact-2-RFS to be useful in other settings. Overall, our study advances the theoretical understanding of several important supertree problems and also provides a new method that should improve scalability of phylogeny estimation methods.

This study suggests several directions for future work. For example, although we showed that Exact-2-RFS produced better RFS criterion scores than FastRFS when used in divide-and-conquer species tree estimation (and similarly GreedyRFS was better than FastRFS for small numbers of source trees in supertree estimation), additional studies are needed to explore its performance, including additional datasets (both simulated and biological datasets) and other leading supertree methods. Similarly, other heuristics using Exact-2-RFS besides GreedyRFS should be developed and studied.

## Acknowledgments

This research was supported by NSF grants 1458652, 1513629, and 1535977 to TW, by the NSF Graduate Research Fellowship DGE-1144245 to EKM, and by the Ira and Debra Cohen Fellowship in Computer Science at the University of Illinois to SAC and EKM. This research is part of the Blue Waters sustained-petascale computing project, which is supported by the National Science Foundation (awards OCI-0725070 and ACI-1238993) and the state of Illinois.

## 6 Notation

### 6.1 Standard Notation

– [*k*] := {1, 2, *…, k*}: the set of integers from 1 to *k*
– *A*:= {*T*_1_, *T*_2_, *…, T*_*N*_} : a profile, i.e., a set of unrooted source trees
– *N* : the number of source trees in a profile
– *T*_*i*_: a source tree in a profile for any *i* ∈ [*N*]
– *V* (*T*), *E*(*T*), *L*(*T*): vertex, edge, and leaf set of a tree *T*
– 𝒯_*S*_ : the set of trees *T* such that *L*(*T*) = *S*
– 𝒯_*S*_^*B*^ : the set of binary trees *T* such that *L*(*T*) = *S*
– *N*_*T*_ (*v*): the set of neighbors of *v* in *T*
– *π*_*e*_: the bipartition of *L*(*T*) induced by deleting an edge *e* from a tree *T*
– *C*(*T*) := {*π*_*e*_ | *e* ∈ *E*(*T*)}: the set of bipartitions that defines a tree *T*
– *T*|_*R*_: the subtree of a tree *T* induced on leaf set *R ⊆ L*(*T*), with degree-two vertices suppressed
– [*A*|*B*]: a bipartition of the set *A* ∪ *B*
– *π*|_*R*_: the bipartition restricted to *R* for *π* = [*A*|*B*], which is [*A ∩ R*|*B ∩ R*]

### 6.2 Notation for RFS and SFS

– RF(*T, T ′*): the Robinson-Foulds (RF) distance between trees *T* and *T ′* (not necessarily having the same leaf set), calculated by |*C*(*T*|_*X*_) *C*(*T ′*|_*X*_) | + |*C*(*T ′*|_*X*_) *C*(*T*|_*X*_)|, where *X* is the shared leaf set.
– SF(*T, T ′*): the split support of trees *T* and *T ′* (not necessarily having the same leaf set), calculated by |*C*(*T* |_*X*_) *∩ C*(*T ′*|_*X*_)|, where *X* is the shared leaf set.
– RF(*T, 𝒜*): the Robinson-Foulds (RF) score of tree *T* with respect to a profile 𝒜, calculated by ∑_*i*∈[*N*]_ RF(*T, T*_*i*_).

SF(*T*, 𝒜): the split support score of tree *T* with respect to a profile 𝒜, calculated by ∑_*i*∈[*N*]_ SF(*T, T*_*i*_).

### 6.3 Notation for Exact-RFS-2

We assume *T*_1_ and *T*_2_ are two binary trees with leaf set *S*_1_ and *S*_2_.

– *X* = *S*_1_ ∩ *S*_2_: the shared leaf set
– *S* = *S*_1_ ∪ *S*_2_: the union of leaf set
– *Π* = 2^*X*^ : the set of all bipartitions of *X*
– *C*(*T*_1_, *T*_2_, *X*) := *C*(*T*_1_|_*X*_) ∪ *C*(*T*_2_|_*X*_): the set of bipartitions of *X* in the backbone trees of *T*_1_ and *T*_2_
– *P* (*e*): the path in *T*_*i*_ from which a backbone edge *e* ∈ *T*_*i*_|_*X*_ is obtained by suppressing degree-two vertices
– *w*(*e*) = |*P* (*e*)|: the weight of edge *e* ∈ *E*(*T*_*i*_|_*X*_)
– *e*_*i*_(*π*): theLedge in *T*_*i*_ that induces the bipartition *π* ∈ *C*(*T*_1_, *T*_2_, *X*)
– *w*^*\**^(*π*) :=∑ _*i*∈[2]_ *w*(*e*_*i*_(*π*)): the weight of bipartition *π* ∈ *Π*, which is the sum of weights of edges in *T*_*i*_|_*X*_ that induces *π*
– *T*_*i*_ *− T*_*i*_|_*X*_ : the subgraph obtained by deleting all vertices and edges of the subgraph of *T*_*i*_ induced on *X*
– Extra(*T*_*i*_) := {*t* | *t* is a component in *T*_*i*_ *− T*_*i*_|_*X*_}: the set of extra subtrees of *T*_*i*_
– *r*(*t*): the root of extra subtree *t*, which is the unique vertex in *V* (*t*) that is adjacent to a vertex in the backbone tree
– 𝒯 ℛ (*e*): the set of extra subtrees attached to *e* ∈ *T*_*i*_|_*X*_, i.e., the set of extra subtrees whose roots are adjacent to internal vertices of *P* (*e*)
– 𝒯 ℛ^*\**^(*π*) := _*i*∈[2]_ 𝒯 ℛ(*e*_*i*_(*π*)): the set of extra subtrees that are attached to edges in *T*_*i*_|_*X*_ that induce *π*
– ℬ𝒫_*i*_(*Q*) := {[*A*|*B*] ∈ *C*(*T*_*i*_|_*X*_) | *A* ⊊ *Q* or *B* ⊊ *Q*}: the set of bipartitions in *C*(*T*_*i*_|_*X*_) with one side being a strict subset of *Q*
– ℬ𝒫 (*Q*) := ℬ𝒫 _1_(*Q*) ∪ *ℬ𝒫* _2_(*Q*): the set of bipartitions in *C*(*T*_1_, *T*_2_, *X*) with one side being a strict subset of *Q*
– 𝒯 ℛ 𝒮_*i*_(*Q*) := ∪_*π∈ℬ𝒫*_*i*_(*Q*)_ 𝒯 ℛ(*e*_*i*_(*π*)): the set of extra subtrees attached to the edges in *T*_*i*_|_*X*_ inducing bipartitions in ℬ𝒫_*i*_(*Q*)
– 𝒯 ℛ 𝒮(*Q*) := 𝒯 ℛ 𝒮_1_(*Q*) ∪ *𝒯 ℛS*_2_(*Q*): the set of extra subtrees attached to edges in *T*_1_|_*X*_ and *T*_2_|_*X*_ inducing bipartitions in ℬ𝒫(*Q*), equivalent to ∪_*π∈ℬ𝒫*(*Q*)_ *𝒯 ℛ\** (*π*)
– 𝒞 = *C*(*T*_1_) ∪ *C*(*T*_2_): the set of bipartitions from input trees *T*_1_ and
– *Π*_*X*_ := {[*A*|*B*] ∈|*C A ∩ X* = *∅* and *B ∩ X* = *∅*}: the set of bipartitions from input trees that are induced by edges on paths connecting vertices of *X*
– *Π*_*Y*_ := {[*A*|*B*] ∈ *C*| *A ∩ X* = *∅* or *B ∩ X* = *∅*}: the set of bipartitions from input trees that are induced by edges not on paths connecting vertices of *X*, i.e., edges in an extra subtree or connecting an extra subtree to the backbone trees
– *p*_*X*_ (*T*) := _*i*∈[2]_ |*C*(*T* |_*S*_*i*) ∩ *C*(*T*_*i*_) ∩ *Π*_*X*_ |: the split support score of *T* contributed by bipartitions in *Π*_*X*_
– *p*_*Y*_ (*T*) := _*i*∈[2]_ |*C*(*T* |_*S*_*i*) ∩ *C*(*T*_*i*_) ∩ *Π*_*Y*_ |: the split support score of *T* contributed by bipartitions in *Π*_*Y*_

## 7 General Theorems and Lemmas on Trees and Bipartitions

The following theorem and corollary gives alternative characterizations of compatibility between two bipartitions.

### Theorem 3

*[21]* **A pair of bipartitions [*A***|***B*] *and* [*A′***|***B′*] *of the same set is compatible if and only if at least one of the four pairwise intersections A ∩ A′, A ∩ B′, B ∩ A′, B ∩ B′ is empty***.

### Corollary 2

*A pair of bipartitions* [*A*|*B*] *and* [*A′*|*B′*] *on the same leaf set is compatible if and only if one side of* [*A*|*B*] *is a subset of one side of* [*A′*|*B′*].

We now provide a lemma and corollary that formalize the relationship between two distinct, yet closely related entities: bipartitions from a tree on leaf set *R ⊆ S* and bipartitions restricted to *R* from a tree on leaf set *S*.

### Lemma 8

*Let T* ∈ *𝒯*_*S*_ *and let π* = [*A*|*B*] ∈ *C*(*T*) *be a bipartition induced by e* ∈ *E*(*T*). *Let R ⊆ S*.

1. – *If R ∩ A* = *∅ or R ∩ B* = *∅, then e ∉ P* (*e′*) *for any e′* ∈ *E*(*T* |_*R*_).
2. – *If R∩A ≠ ∅ and R∩B ≠ ∅, then for any π′* ∈ *C*(*T* |_*R*_) *induced by e′* ∈ *E*(*T* |_*R*_), *π*|_*R*_ = *π′ if and only if e* ∈ *P* (*e′*).

*Proof*. Let *T*_*R*_ be the minimal subtree of *T* that spans *R*. It follows that the leaf set of *T*_*R*_ is *R* and *T*|_*R*_ is obtained from *T*_*R*_ by suppressing all degree-two vertices.

(Proof of 1) We first claim that if *R ∩ A* = *∅* or *R ∩ B* = *∅*, then *e ∉ E*(*T*_*R*_). Assume by way of contradiction that *e* ∈ *E*(*T*_*R*_). There are then two non-empty components in *T*_*R*_ *− e*. Since *e* induces [*A*|*B*] in *T*, the two components in *T*_*R*_ *− e* have leaf set *R ∩ A* and *R ∩ B*, which contradicts the fact that one intersection is empty. Therefore, *e ∉ E*(*T*_*R*_). Furthermore, every edge *e′* ∈ *E*(*T* _*R*_) comes from a path in *T*_*R*_. Since *e ∉ E*(*T*_*R*_), then *e / P* (*e′*) for any *e′* ∈ (*T* _*R*_).

(Proof of 2) If *R ∩ A* ≠ *∅* and *R ∩ B* ≠ *∅*, then *e* is required to connect *R ∩ A* with *R ∩ B* in *T* (since *e* connects *A* with *B*). Thus, *e* is in any subtree of *T* spanning *R*; in particular, *e ∈ E*(*T*_*R*_). Fix any *π′* ∈ *C*(*T*|_*R*_) induced by *e′* ∈ (*T*|_*R*_). Note that the bipartition induced by *P* (*e′*) in *T*_*R*_ equals the bipartition induced by *e′* in *T*|_*R*_, i.e., *π′*. For one direction of the proof, suppose *e* ∈ *P* (*e′*). Because internal vertices of *P* (*e′*) in *T*_*R*_ do not connect to any leaves, the bipartition induced by the path *P* (*e′*) in *T*_*R*_ equals the bipartition induced by any of its edges (in particular, *e*). Since *e* induces [*A*|*B*] in *T*, it induces [*R ∩ A*|*R ∩ B*] in *T*_*R*_. Then *π′* = [*R ∩ A*|*R ∩ B*] = *π*|_*R*_. On the other hand, if *π*|_*R*_ = *π′*, then *π′* induces [*R ∩ A*|*R ∩ B*] in *T* |_*R*_. It follows that *P* (*e′*) also induces [*R ∩ A*|*R ∩ B*] in *T*_*R*_. Suppose *e* ∈ *P* (*e*^*\**^) for some edge *e*^*\**^ ∈ *E*(*T* |_*R*_) such that *e*^*\**^ *≠ e′*. Then, by the previous argument, *π*_*e*\*_ = [*R* ⋃ *A*|*R* ⋃ *B*], which contradicts the fact that *e*^*\**^ and *e′* are different edges. Therefore, *e* ∈ *P* (*e′*).

The next corollary follows easily from Lemma 8.

### Corollary 3

*Let T be a tree with leaf set S and let π* = [*A*|*B*] ∈ *C*(*T*) *be a bipartition induced by e* ∈ *E*(*T*). *Let R ⊆ S such that R ∩ A ≠ ∅ and R ∩ B ≠∅. Then π*|_*R*_ ∈ *C*(*T* |_*R*_).

In the following lemma, we characterize the vertex that we can split to add a compatible bipartition into a tree. An example can be seen in Figure 6.

**Fig. 6:**
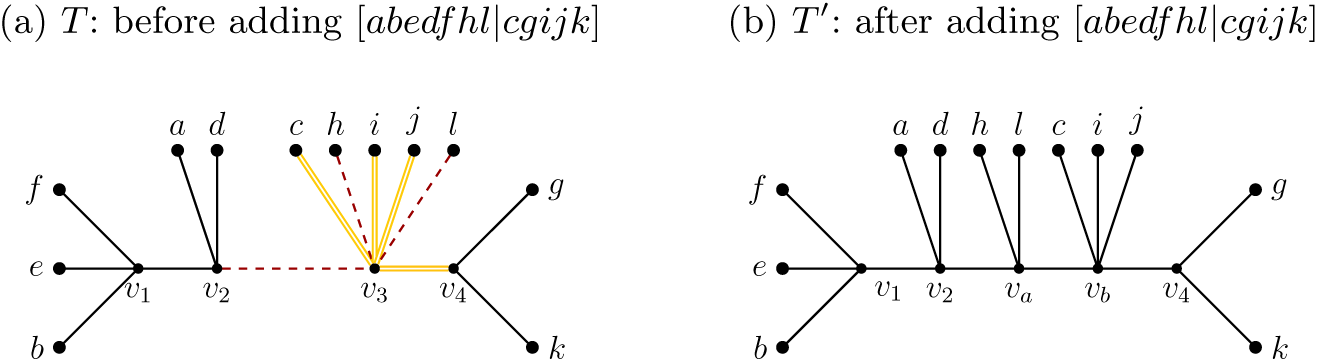
Splitting a vertex in a tree *T* to add a compatible bipartition [*A*|*B*] = [*abedf hl*|*cgijk*]. The vertex *v*_3_ satisfies the requirement that no component in *T − v*_3_ has leaves from both *A* and *B*. Let *N*_*A*_ (*N*_*B*_) denote the neighbors of *v*_3_ that are in a component containing a leaf in *A* (*B*) in *T −v*_3_. Then *N*_*A*_ = {*v*_2_, *h, l*} and *N*_*B*_ = {*c, i, j, v*_4_}. We split *v*_3_ into *v*_*a*_ and *v*_*b*_. We then make *N*_*A*_ the neighbors of *v*_*a*_, and *N*_*B*_ the neighbors of *v*_*b*_. Then (*v*_*a*_, *v*_*v*_) induces [*abedf hl*|*cgijk*] in *T ′*.

### Lemma 9

*Let T be a tree with leaf set S. Let π* = [*A*|*B*] *be a bipartition of S such that π ∉ C*(*T*), *but π is compatible with C*(*T*). *Then there exists a unique vertex v* ∈ *V* (*T*) *such that no component of T − v has leaves from both A and B. Furthermore, we can split the neighbors of v, N*_*T*_ (*v*), *into two sets N*_*A*_ *and N*_*B*_, *where N*_*A*_ *contains neighbors whose corresponding components contain a leaf in A and N*_*B*_ *contains neighbors whose corresponding components contain a leaf from B. By replacing v with two vertices v*_*a*_ *and v*_*b*_, *making v*_*a*_ *adjacent to all the vertices in N*_*A*_ *and v*_*b*_ *adjacent to all the vertices in N*_*B*_, *and then adding an edge between v*_*a*_ *and v*_*b*_, *we create a tree T ′ such that C*(*T ′*) = *C*(*T*) ∪ *{π*}.

*Proof*. By definition of compatibility, there exists a tree *T ′* such that *C*(*T ′*) = *C*(*T*) ∪*{π*}. Let *e* = (*v*_*a*_, *v*_*b*_) be the edge that induces *π* in *T ′* such that the component containing *v*_*a*_ in *T*′ *−* (*v*_*a*_, *v*_*b*_) has leaf set *A* and the component containing *v*_*b*_ in *T*′ *−* (*v*_*a*_, *v*_*b*_) has leaf set *B*. Since *π ∉ C*(*T*), when we contract (*v*_*a*_, *v*_*b*_), then *T ′* becomes *T*. Let *v* be the vertex of *T* corresponding to the vertex of *T ′* created from contracting (*v*_*a*_, *v*_*b*_). Let *N*_*a*_, *N*_*b*_ be the neighbors of *v*_*a*_ and *v*_*b*_ in *T ′ −* (*v*_*a*_, *v*_*b*_), respectively. Let *N*_*A*_, *N*_*B*_ be vertices in *T* corresponding to *N*_*a*_ and *N*_*b*_. We note that *N*_*A*_ ∪ *N*_*B*_ = *N*_*T*_ (*v*). Since in *T ′ −* (*v*_*a*_, *v*_*b*_), no vertex in *N*_*a*_ can reach any vertex of *B*, the same is true in *T ′ − v*_*a*_ *− v*_*b*_. Since *v*_*a*_ is in the component of *A* in *T ′ −* (*v*_*a*_, *v*_*b*_), so are all vertices of *N*_*a*_. Then each vertex in *N*_*a*_ must be able to reach some vertex of *A* in *T ′ − v*_*a*_ *− v*_*b*_ by either being a leaf in *A* or in the same component of some leaf in *A*. Similarly, in *T ′ − v*_*a*_ *− v*_*b*_, no vertex of *N*_*b*_ can reach any vertex of *A*, but every vertex of *N*_*b*_ can reach some vertex of *B*. By construction, *T ′ − v*_*a*_ *− v*_*b*_ is identical to *T − v*, and thus *N*_*A*_ (and *N*_*B*_ respectively) is a set of neighbors of *v* that can reach some vertex of *A* (*B*) but no vertex of *B* (*A*). Therefore, *v* is the vertex desired.

To obtain *T ′* from *T*, we can replace *v* by two new vertices *v*_*a*_, *v*_*b*_ with an edge between them. We also connect all vertices in *N*_*A*_ to *v*_*a*_ and all vertices in *N*_*B*_ to *v*_*b*_. Then it is easy to see that (*v*_*a*_, *v*_*b*_) induces *π* in *T ′*. □

## 8 Proofs for Section 3

### Lemma 1

*Given an input set 𝒜 of source trees, a tree* 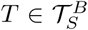 *is an optimal solution for* RFS*(𝒜) if and only if it is an optimal solution for* SFS(𝒜).

*Proof*. Let *N ≥* 2 be any integer. Let *T*_1_, *T*_2_, *…, T*_*N*_ and *S*_1_, *S*_2_, *…, S*_*N*_ be defined as from problem statement of RFS. Let *T* be any binary tree of leaf set *S*. Then 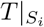 is also binary and thus 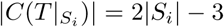. For any *i* ∈ [*N*], we have

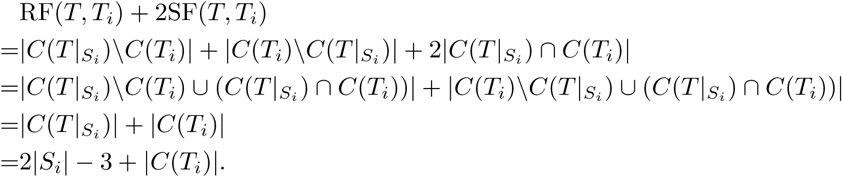

Taking the sum of the equations, we have

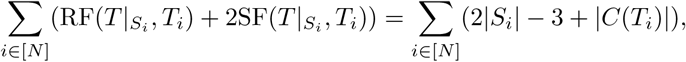

which is a constant. Therefore, for any binary tree *T* and any profile *A* of source trees, the sum of *T* ‘s RFS score and twice *T* ‘s split support score is the same, independent of *T*. This implies that minimizing the RFS score is the same as maximizing the split support score. Although this argument depends on the output tree being binary, it does not depend on the input trees being binary. Hence, we conclude that RFS and SFS have the same set of optimal supertrees. □

### Lemma 2

*Proof*. Since *T*_1_ and *T*_2_ have different leaf sets, *C*(*T*_1_) and *C*(*T*_2_) are disjoint. Since *Π*_*Y*_ ⊆ *C*(*T*_1_) ∪ *C*(*T*_2_), *C*(*T*_1_) ∩ *Π*_*Y*_ and *C*(*T*_2_) ∩ *Π*_*Y*_ form a disjoint decomposition of *Π*_*Y*_. By definition of *p*_*Y*_ (·), for any tree *T* of leaf set *S*,

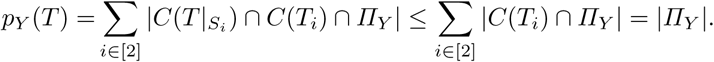

Fix any *π* = [*A*|*B*] ∈ *Π*_*Y*_. Suppose *π* ∈ *C*(*T*_*i*_) and is induced by *e* ∈ *E*(*T*_*i*_) for some *i* ∈ [2]. By definition of *Π*_*Y*_, either *A ∩ X* = *∅* or *B ∩ X* = *∅*. By Lemma 8, *e ∉ P* (*e′*) for any backbone edge *e′* ∈ *E*(*T*_*i*_|_*X*_). Therefore, either *e* is an internal edge in an extra subtree in Extra(*T*_*i*_), or *e* connects one extra subtree in Extra(*T*_*i*_) to the backbone tree. In either case, the construction of *T*_init_ ensures that *e* is also present in *T*_init_|_*S*_*i* and thus *π* ∈ *C*(*T*_init_|_*S*_*i*). Therefore, each bipartition *π* ∈ *Π*_*Y*_ contributes 1 to |*C*(*T*_init_|_*S*_*i*) ∩ *C*(*T*_*i*_) ∩ *Π*_*Y*_ | for exactly one index *i* ∈ [2] and thus it contributes 1 to *p*_*Y*_ (*T*_init_). Hence, *p*_*Y*_ (*T*_init_) = |*Π*_*Y*_ |. □

### Lemma 3

*Let π* = [*A*|*B*] ∈ *Π. Let T* ∈ *𝒯*_*S*_ *be any tree with leaf set S such that π ∉ C*(*T* |_*X*_) *but π is compatible with C*(*T* |_*X*_). *Let T ′ be a refinement of T such that for all* 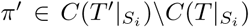 *for some i* ∈ [2], *π′*|_*X*_ = *π. Then, p*_*X*_ (*T ′*) *− p*_*X*_ (*T*) *≤ w*^*\**^(*π*).

*Proof*. By definition of *p*_*X*_ (·),

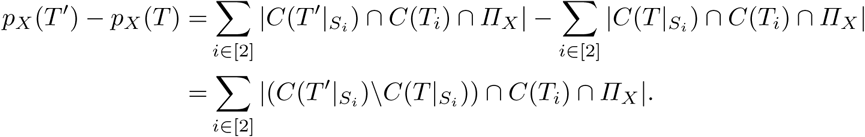

Therefore, we only need to prove that

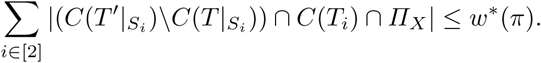

We perform a case analysis, as follows: Case (1): *π ∉ C*(*T*_1_, *T*_2_, *X*), Case (2): *π* ∈ *C*(*T*_1_|_*X*_)*ΔC*(*T*_2_|_*X*_) = (*C*(*T*_1_|_*X*_) *\ C*(*T*_2_|_*X*_)) ∪ (*C*(*T*_2_|_*X*_) *\ C*(*T*_1_|_*X*_)), and Case (3): *π* ∈ *C*(*T*_1_|_*X*_) ∩ *C*(*T*_2_|_*X*_).

Case (1): Let *π ∉ C*(*T*_1_, *T*_2_, *X*). We recall that *w* (*π*) = 0. Assume by way of contradiction that there exists a bipartition 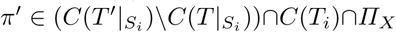 for some *i* ∈ [2]. Since *π ∉ C*(*T*_1_, *T*_2_, *X*) and *π′*|_*X*_ = *π*, by Corollary 3, *π′ ∉ C*(*T*_*i*_) for any *i* ∈ [2]. This contradicts the fact that *π′* ∈ *C*(*T*_*i*_) for some *i* ∈ [2]. Therefore, the assumption that there exists such a bipartition *π′* is wrong and 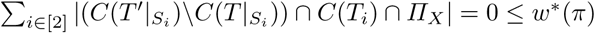.

Case (2): Let *π* ∈ *C*(*T*_1_|_*X*_)*ΔC*(*T*_2_|_*X*_). Assume without loss of generality that *π* ∈ *C*(*T*_1_|_*X*_)*\C*(*T*_2_|_*X*_). Then, we have *w*^*\**^(*π*) = *w*(*e*_1_(*π*)). Let 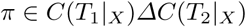 for some *i* ∈ [2]. Then we have *π′*|_*X*_ = *π* by assumption of the lemma. Since *π ∉ C*(*T*_2_|_*X*_), by Corollary 3, we have *π′ ∉ C*(*T*_2_) and thus *π′* ∈ *C*(*T*_1_). By Lemma 8, the edge which induces *π′* in *T*_1_ is an edge on *P* (*e*_1_(*π*)). Since there are *w*(*e*_1_(*π*)) edges on *P* (*e*_1_(*π*)), there are at most *w*(*e*_1_(*π*)) distinct bipartitions *π′*, proving the claim.

Case (3): Let *π* ∈ *C*(*T*_1_|_*X*_) ∩ *C*(*T*_2_|_*X*_). Then we have *w*^*\**^(*π*) = *w*(*e*_1_(*π*)) + *w*(*e*_2_(*π*)). Fix any *π′* ∈ *C*(*T*_*i*_) ∩ *Π*_*X*_ ∩ (*C*(*T ′*|_*S*_*i*)*\C*(*T* |_*S*_*i*)) for any *i* ∈ [2]. Since *π′* ∈ *C*(*T*_*i*_) and *π′*|_*X*_ = *π* ∈ *C*(*T*_*i*_|_*X*_), by Lemma 8, the edge *e′* that induces *π′* is an edge on *P* (*e*_*i*_(*π*)). Since there are *w*(*e*_*i*_(*π*)) edges on *P* (*e*_*i*_(*π*)), there are at most *w*(*e*_*i*_(*π*)) distinct bipartitions *π′* in 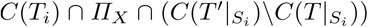. Therefore,

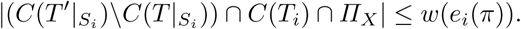

Hence,

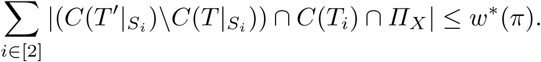

□

### Lemma 4

*For any compatible set F ⊆ Π, let T* ∈ 𝒯_*S*_ *be any tree with leaf set S such that C*(*T* |_*X*_) = *F*. *Then p*_*X*_ (*T*) *≤ w*^*\**^(*F*).

*Proof*. Fix an arbitrary ordering of bipartitions in *F* and let them be *π*_1_, *π*_2_, *…, π*_*k*_, where *k* = |*F* |. Let *F*_*j*_ = *{π*_1_, *…, π*_*j*_} for any *j* ∈ *{*0, 1, *…, k*}. In particular, *F*_0_ = *∅* and *F*_*k*_ = *F*. Let *T*^*j*^ be obtained by contracting all edges in *P* (*e*) for any *e* ∈ *E*(*T* |_*X*_) such that *π*_*e*_ *∉ F*_*j*_. Then, *C*(*T*^*j*^|_*X*_) = *F*_*j*_. For each *j* ∈ [*k*], *C*(*T*^*j*^|_*X*_)*\C*(*T*^*j−*1^|_*X*_) = *{π*_*j*_}. Fix *j* ∈ [*k*] and fix any 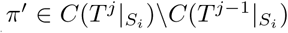 for some *i* ∈ [2]. By Lemma 8, we have *π′*|_*X*_ ∈ *C*(*T*^*j*^|_*X*_). We also know *π′*|_*X*_ *∉ C*(*T*^*j−*1^|_*X*_) as otherwise 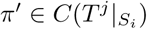 by construction, which is a contradiction. Then *π′*|_*X*_ ∈ *C*(*T*^*j*^|_*X*_)*\C*(*T*^*j−*1^|_*X*_) = *{π*_*j*_}. Therefore, for any *j* ∈ [*k*], *T*^*j*^ is a refinement of *T*^*j−*1^ such that for any 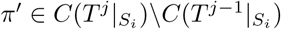 for some *i* ∈ [2], *π′*|_*X*_ = *π*_*j*_. Hence we can apply Lemma 3 and we have *p*_*X*_ (*T* ^*j*^) − *p*_*X*_ (*T* ^*j−*1^) *w*^*\**^(*π*_*j*_). Therefore, by telescoping sum,

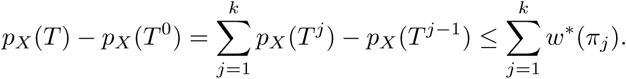

Since *C*(*T* ^0^|_*X*_) = *∅*, by Corollary 3, *C*(*T* ^0^|_*S*_) ∩ *Π*_*X*_ = *∅* for both *i* ∈ [2]. Then, 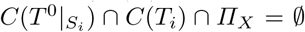 for both *i* ∈ [2], which implies *p*_*X*_ (*T* ^0^) = 0. Thus, *p*_*X*_ (*T*) *≤ ∑*_*π∈F*_ *w*^*\**^(*π*), as desired.

Let *T*_init_ be the tree defined in Algorithm 1. We have the following proposition about *p*_*X*_ (*T*_init_), which is needed for the proof of Proposition 1.

### Proposition 3

*p*_*X*_ (*T*_init_) = 2|*X*|.

*Proof*. For each *v ∈ X*, consider the bipartition *π*_*v*_ = [*{v*} |*S\ {v*}] of *T*_init_ induced by the edge that connects the leaf *v* to the center 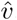. It is easy to see that *π*_*v*_|_*S*_*i* = [*{v*} | *S*_*i*_*\{v*}] ∈ *C*(*T*_*i*_) for any *i* ∈ [2] as *π*_*v*_|_*S*_*i* is a trivial bipartition of *S*_*i*_. By construction, we also have *π*_*v*_|_*S*_*i* ∈ *C*(*T*_init_|_*S*_*i*). We also know *π*_*v*_|_*S*_*i* ∈ *Π*_*X*_ as both sides of *π*_*v*_ have non-empty intersections with *X*. Thus, *π*_*v*_|_*S*_*i* ∈ *C*(*T*_init_|_*S*_*i*) ∩ *C*(*T*_*i*_) ∩ *Π*_*X*_ for any *i* ∈ [2]. So for each *v* ∈ *X, π*_*v*_|_*S*_1 and *π*_*v*_|_*S*_2 each contributes 1 to *p*_*X*_ (*T*_init_). Therefore, *p*_*X*_ (*T*_init_) *≥* 2|*X*|.

Fix any bipartition *π* = [*A*|*B*] induced by any other edge 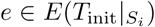 for any *i* ∈ [2]. By construction of *T*_init_, *e* must be an edge in an extra subtree or connecting an extra subtree to the center 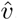, i.e., one component in *T − e* does contain any leaf of *X*. Therefore, either *A ⊆ S\X* or *B ⊆ S\X*, which implies 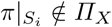 for any *i* ∈ [2]. Hence, there is no other bipartition of *T*_init_ such that when restrict to *S*_*i*_ contributes to *p*_*X*_ (*T*_init_). Therefore, *p*_*X*_ (*T*_init_) = 2|*X*|.

### Proposition 1

*Let* 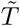 *be the tree constructed after line 11 of Algorithm 1, then* 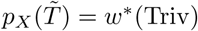.

*Proof*. Let *π* = [{*a*}|*B*] be a trivial bipartition of *X*. We know both *e*_1_(*π*) and *e*_2_(*π*) exist, and we abbreviate them with *e*_1_ and *e*_2_. We number the extra subtrees in *T ℛ* (*e*_1_) as *t*_1_, *t*_2_, *…, t*_*p*_, where *p* = *w*(*e*_1_) *−* 1, such that *t*_1_ is the closest to *a* in *T*_1_. Similarly, we number extra subtrees in (*e*_2_) as 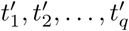, where *q* = *w*(*e*_2_) *−* 1, such that *t′*_1_ is the closest to *a* in *T*_2_. For each *k* ∈ [*w*(*e*_1_)], we define

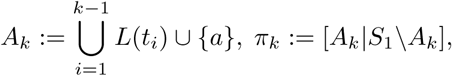

and for each *k* ∈ [*w*(*e*_2_)], we define

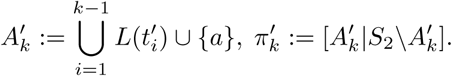

It follows by definition that *π*_*k*_ for any *k* ∈ [*w*(*e*_1_)] is the bipartition induced by the *k*th edge on *P* (*e*_1_) in *T*_1_, where the edges are numbered starting from the side of *a*. This implies *π*_*k*_ ∈ *C*(*T*_1_) for any *k* ∈ [*w*(*e*_1_)]. Similarly, 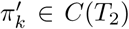 for any *k* ∈ [*w*(*e*_2_)]. In particular, we notice that *π*_1_ = [{*a*}|*S*_1_*\{a*}] and 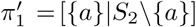. Clearly, all these bipartitions (*π*_*k*_ and 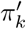 for any *k*) are in *Π*_*X*_ because both sides have none empty intersection with *X*.

Recall that Algorithm 1 moves all extra subtrees in *T ℛ*^*\**^(*π*) onto the edge 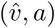 and orders them in a way to add our desired bipartitions. In particular, the extra subtrees are ordered such that subtrees from *T ℛ* (*e*_1_) and subtrees from *T ℛ* (*e*_2_) are side by side and the attachments of *T ℛ* (*e*_*i*_) match their attachment on *e*_*i*_ exactly (i.e., *t*_1_ or 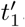, respectively, is closest to *a*). It is easy to see that as a result of such ordering of the extra subtrees, we have 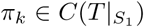) for any *k* ∈ [*w*(*e*_1_)] and 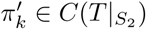 for any *k* ∈ [*w*(*e*_2_)], where *T* is the tree obtained after adding *π* to the backbone through line 8 of Algorithm 1. Therefore, the algorithm increases 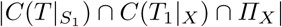 by *w*(*e*_1_) *−* 1, because 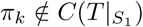 before the step for all *k* ∈ [*w*(*e*_1_)] except *k* = 1 (since 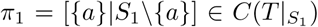). Similarly, the algorithm increases 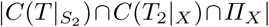 by *w*(*e*_2_) *−* 1. Overall *p*_*X*_ (*T*) is increased by *w*(*e*_1_) + *w*(*e*_2_) *−* 2 = *w*^*\**^(*π*) *−* 2 by running one execution of line 8 in Algorithm 1 on *T* and *π*.

It is easy to see that line 8 of Algorithm 1 never destroys bipartitions of *S*_1_ or *S*_2_ already in *T*, so we have

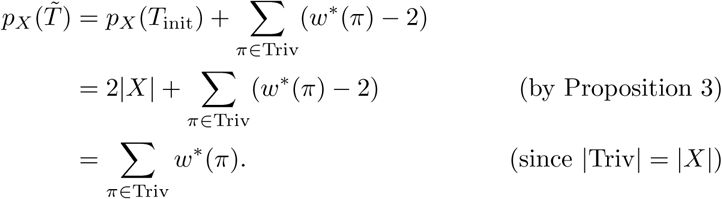

Lemma 10 proves that the auxiliary data structures of Algorithm 1 and 2 are maintaining the desired information and that the algorithm can split the vertex and perform the detaching and reattaching of the extra subtrees correctly. These invariants are important to the proof of Lemma 5.

### Lemma 10

*At any stage of the Algorithm 1 after line* 12, *we have the following invariants of T and the auxiliary data structures H and sv:*

1. *For any bipartition π* ∈ NonTriv, *sv*(*π*) *is the vertex to split to add π to C*(*T* |_*X*_). *For any internal vertex v, the set of bipartitions H*(*v*) ⊆ NonTriv *is the set of bipartitions which can be added to C*(*T* |_*X*_) *by splitting v*.
2. *For any π* = [*A*|*B*] ∈ *H*(*v*), *for all t* ∈ *T ℛ\**(*π*), *the root of t is a neighbor of v*.
3. *For any π* = [*A*|*B*] ∈ *C*(*T* |_*X*_) *induced by edge e, let C*(*A*), *C*(*B*) *be the components containing the leaves of A and B in T* |_*X*_ *− e. Then*,
  a. *all t* ∈ *𝒯 ℛS*(*A*) *are attached to an edge or a vertex in C*(*A*)
  b. *all t* ∈ 𝒯 ℛ 𝒮(*B*) *are attached to an edge or a vertex in C*(*B*).

*Proof*. We prove the invariants by induction on the number of refinement steps *k* performed on *T*. When *k* = 0, we have 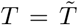 and *T*|_*X*_ is a star with leaf set *X* and center vertex 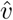. Thus all bipartitions in NonTriv are compatible with *C*(*T* |_*X*_). For any *π* ∈ NonTriv, 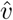 is the vertex to refine in *T* |_*X*_ to add *π* to *C*(*T* |_*X*_). Therefore, it is correct that 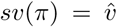 for every *π* ∈ NonTriv and 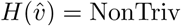. The roots of all extra subtrees in *𝒯 ℛ\**(*π*) for any *π* ∈ NonTriv are all neighbors of 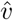, so invariant 2 also holds. For any *π* ∈ *C*(*T* |_*X*_) = Triv, let *π* = [{*a*}|*B*]. It is easy to see that since *a* is a leaf, 𝒯 ℛ 𝒮_*i*_({*a*}) = *∅* and 𝒯 ℛ 𝒮_*i*_(*B*) = Extra(*T*_*i*_)*\𝒯 ℛ* (*e*_*i*_(*π*)) for both *i* ∈ [2]. Then *𝒯 ℛS*({*a*}) = *∅* and *𝒯 ℛS*(*B*) = (Extra(*T*_1_) ∪ Extra(*T*_2_))*\𝒯 ℛ\**(*π*). Therefore, invariant 3(a) trivially holds as *𝒯 ℛS*({*a*}) = *∅*. Since *C*({*a*}) is the vertex *a* and *C*(*B*) is the rest of the star of *T*|_*X*_, all *t* ∈ *𝒯 ℛS* (*B*) are attached to an edge or a vertex in *C*(*B*), then invariant 3(b) holds. This proves invariant 3 and thus concludes our proof for the base case.

Assume that all invariants hold after any *k′ < k* steps of refinement. Let *π* = [*A*|*B*] be the bipartition to add in the *k*th refinement step. We will show that after the *k*th refinement step, i.e., one execution of Algorithm 2, the invariants still hold for the resulting tree *T ′*. Since *v* = *sv*(*π*) at the beginning of Algorithm 2, *π* can be added to *C*(*T* |_*X*_) by splitting *v*. By Lemma 9, there exists a division of neighbors of *v* in *T* |_*X*_ into *N*_*A*_ ∪*N*_*B*_ such that *N*_*A*_ (or *N*_*B*_ respectively) consists of neighbors of *v* which can reach vertices of *A* (or *B*) but not *B* (or *A*) in *T* |_*X*_ *−v*. Then, the algorithm correctly finds *N*_*A*_ and *N*_*B*_ and connects *N*_*A*_ to *v*_*a*_ and *N*_*B*_ to *v*_*b*_ so the new edge (*v*_*a*_, *v*_*b*_) induces the bipartition *π* =∈ [*A*|*B*] in *T*|_*X*_. For any vertex *u* other than *v* and any bipartition *π′* ∈ *H*(*u*), the invariants 1 and 2 still hold after Algorithm 2 as we do not change *H*(*u*), *sv*(*π′*), or the extra subtrees attached to *u*. For any bipartition *π′* = *H*(*v*) such that *π′* ≠ *π*, if *π′* is not compatible with *π*, then it cannot be added to *C*(*T ′*|_*X*_) since *π* is added, so the algorithm correctly discards *π′* and does not add it to *H*(*v*_*a*_) or *H*(*v*_*b*_). If *π′* is compatible with *π*, we will show that the invariants 1 and 2 still hold for *π′*.

Fix any *π′* = [*A′*|*B′*] *H*(*v*) s.t. *π′* = *π* and *π′* is compatible with *π*. By Corollary 2, one of *A′* and *B′* is a subset of one side of [*A B*]. Assume without loss of generality that *A′ ⊆ A* (other cases are symmetric). Then we have *B ⊆ B′*. In this case, Algorithm 2 adds *π′* to *H*(*v*_*a*_) and set *sv*(*π*) = *v*_*a*_. We will show that this step preserves the invariants. Since *π′* ∈ *H*(*v*), before adding *π* we can split *v* to add *π′* to *C*(*T* |_*X*_). Then there exists a division of neighbors of *v* in *T* |_*X*_ into *N*_*A*_*t* and *N*_*B*_*t* such that *N*_*A*_*t* (or *N*_*B*_*t*, respectively) consists of neighbors of *v* which can reach vertices of *A′* (or *B′*) in *T* |_*X*_ *− v*. It is easy to see that *N*_*A*_*t ⊆ N*_*A*_ and *N*_*B*_ ⊆ *N*_*B*_*t*. Since *N*_*A*_ ∪ *N*_*B*_ = *N*_*A*_*t* ∪ *N*_*B*_*t* = *N*_*T* |*X*_ (*v*), we have *N*_*A*_*\N*_*A*_*t* = *N*_*B*_*t \N*_*B*_. Since all vertices in *N*_*B*_ are connected to *v*_*b*_ in *T ′* while vertices in *N*_*B*_*t \N*_*B*_ are connected to *v*_*a*_, *N*_*B*_*t \N*_*B*_ ∪ {*v*_*b*_} is the set of all neighbors of *v*_*a*_ which can reach leaves of *B′* in *T ′*|_*X*_ *− v*_*a*_. Then *N*_*T*_ *t*|_*X*_ (*v*_*a*_) = *N*_*A*_ ∪ {*v*_*b*_} = *N*_*A*_*t ∪* (*N*_*A*_*\N*_*A*_*t ∪ {v*_*b*_}) = *N*_*A*_*t ∪* (*N*_*B*_*t \N*_*B*_ ∪ *v*_*b*_) implies that *N*_*A*_*t* and *N*_*B*_*t\ N*_*B*_ ∪{*v*_*b*_} gives an division of neighbors of *v*_*a*_ such that *N*_*A*_*t* are the neighbors that can reach leaves of *A′* in *T ′*|_*X*_ *− v*_*a*_ and *N*_*B*_*t \N*_*B*_ ∪{*v*_*b*_} are the neighbors that can reach leaves of *B′* in *T ′*|_*X*_ *− v*_*a*_. Such a division proves that *v*_*a*_ is the correct vertex to refine in *T ′* _*X*_ to add *π′* to *C*(*T ′* _*X*_) after the *k*th refinement. Therefore, invariant 1 holds with respect to *π′*. Since *π′* ∈ *H*(*v*) before adding *π*, we also have for all *t* ∈ *𝒯 ℛ*^*\**^ (*π′*), the root of *t* is a neighbor of *v* before adding *π*. Since *A′ ⊆ A, π′* ∈ (*A*) and thus *𝒯 ℛ* ^*\**^(*π*) ⊆*𝒯 ℛS* (*A*). Then, Algorithm 2 correctly attaches roots of all trees in *𝒯 ℛ* ^*\**^(*π′*) to *v*_*a*_. Therefore invariant 2 holds for *π′*.

We have shown that invariants 1 and 2 hold for the tree *T ′* with the auxiliary data structures *H* and *sv*. Next, we show that invariant 3 holds. Since *π* is the only bipartition in *C*(*T ′* _*X*_) that is not in *C*(*T* _*X*_), we only need to show two things: i) for any *π′ C*(*T* _*X*_), the invariant 3 still holds, ii) invariant 3 holds for *π*. We first show i). Fix *π′* = [*A′ B′*] *C*(*T* _*X*_). Since *π* is compatible with *π′*, by Corollary 2, one of *A′* and *B′* is a subset of one of *A* and *B*. We assume without loss of generality that *A′ A*. Therefore, *B B′*. Let *C*(*A′*), *C*(*B′*) be the components containing the leaves of *A′* and *B′* in *T* _*X*_ *e′*, where *e′* induces *π′*. Since *C*(*A′*) is unchanged after the refinement, invariant 3(a) is trivially true. Since *B B′, C*(*B*) is a subgraph of *C*(*B′*) and *v C*(*B′*). During the refinement, *v* is split into *v*_*a*_ and *v*_*b*_, both of which are still part of *C*(*B′*). Since all *t* ∈ *T ℛS*(*B*) are attached to an edge or a vertex in *C*(*B′*) before refinement and any extra subtree attached to *v* before is now on either *v*_*a*_, or *v*_*b*_, or (*v*_*a*_, *v*_*b*_), they are all still attached to an edge or a vertex in *C*(*B′*). Thus, the invariant 3 holds with respect to *π′*.

For ii), we show invariant 3(a) holds for *π* and 3(b) follows the same argument. For any extra subtree in *t* ∈ *𝒯 ℛS*(*A*), if it was attached to *v* before refinement, then it is now attached to *v*_*a*_, which is in *C*(*A*). If it was not attached to *v* before refinement, then let *N*_*B*_ be as defined from Algorithm 2. For any bipartition *π′* = [*A′*|*B′*] induced by (*v, u*) where *u ∈ N*_*B*_. We know that (*v, u*) ∈ *C*(*B*) and thus either *A′ ⊆ B* or *B′ ⊆ B*. Assume without loss of generality that *B′ ⊆ B*. Then we have ℬ𝒫 (*B′*) ∪ {*π′} ⊆ℬ𝒫* (*B*) and thus 𝒯 ℛ 𝒮 (*B′*) ∪*𝒯 ℛ*^*\**^(*π′*) ⊆𝒯 ℛ 𝒮 (*B*). We note that 𝒯 ℛ 𝒮 (*A*) and *𝒯 ℛS* (*B*) are disjoint. Since *t ∈𝒯 ℛS* (*A*), we know *t ∉𝒯 ℛS* (*B*), then *t ∉ 𝒯 ℛS* (*B′*) ∪ *𝒯 ℛS* ^*\**^(*π′*). Let *C*(*A′*), *C*(*B′*) be the components containing the leaves of *A′* and *B′* in *T*|_*X*_ *−* (*v, u*). Then *C*(*A′*) contains *v* and *C*(*B′*) contains *u*. Since *t ∉ 𝒯 ℛ* ^*\**^(*π′*), it cannot be attached to (*v, u*). Also by the invariant 3 with respect to *π′, t* is not attached any vertex or edge in *C*(*B′*). Since this is true for every neighbor of *v* in *N*_*B*_, *t ∉ C*(*B*) as *C*(*B*) consists of only edges connecting *v* to a neighbor *u* ∈ *N*_*B*_ and the component containing *u*. Since *t* was not attached to *v* before the refinement, *t* is not attached to (*v*_*a*_, *v*_*b*_) or *C*(*B*) after the refinement, then *t* must be attached to some edge or vertex in *C*(*A*). This proves invariant 3(a) for *π* and thus the inductive proof. □

### Lemma 5

*Let T be a supertree computed within Algorithm 1 at line* 14 *immediately before a refinement step. Let π* = [*A*|*B*] ∈ NonTriv ∩ *I. Let T ′ be a refinement of T obtained from running Algorithm 2 with supertree T, bipartition π, and the auxiliary data structures H and sv. Then, p*_*X*_ (*T ′*) *− p*_*X*_ (*T*) = *w*^*\**^(*π*).

*Proof*. Since *I* corresponds to an independent set in the incompatibility graph *G*, all bipartitions in *I* are compatible. Since *C*(*T* |_*X*_) ⊆ Triv ∪ (NonTriv ∩ *I*) = *I, π* ∈ NonTriv ∩ *I* must be compatible with *C*(*T* |_*X*_), then there is a vertex to split to add *π* to *C*(*T* |_*X*_). By invariant 1 of Lemma 10, *v* = *sv*(*π*) is the vertex to split to add *π* to *T* |_*X*_ and thus Algorithm 2 correctly splits *v* into *v*_*a*_ and *v*_*b*_ and connects them to appropriate neighbors such that in *T ′*|_*X*_, (*v*_*a*_, *v*_*b*_) induces *π*.

We abbreviate *e*_1_(*π*) and *e*_2_(*π*) by *e*_1_ and *e*_2_. We number the extra subtrees attached to *e*_1_ as *t*_1_, *t*_2_, *…, t*_*p*_, where *p* = *w*(*e*_1_) *−* 1 and *t*_1_ is the closest to *A* in *T*_1_. Similarly, we number the extra subtrees attached to *e*_2_ as 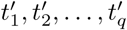, where *q* = *w*(*e*_2_) *−* 1 and 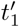 is the closest to *A* in *T*_2_.

For any set *𝒯* of trees, let *L*(*𝒯*) denote the union of the leaf set of trees in *𝒯*. We note that if *e*_*i*_ exists, Extra(*T*_*i*_) = *𝒯 ℛS*_*i*_(*A*) ∪ *𝒯 ℛS*_*i*_(*B*) ∪ *𝒯 ℛ* (*e*_*i*_). Thus, *A ∪ L*(*𝒯 ℛS*_*i*_(*A*)) ∪ *L*(*𝒯 ℛ* (*e*_*i*_)) ∪ *L*(*𝒯 ℛS*_*i*_(*B*)) ∪ *B* = *S*_*i*_ for *i* ∈ [2].

For each *k* ∈ [*w*(*e*_1_)], we define

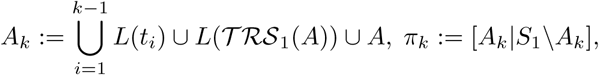

and for each *k* ∈ [*w*(*e*_2_)], we define

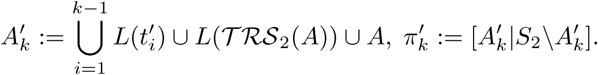

We know that for each *k* ∈ [*w*(*e*_1_)],

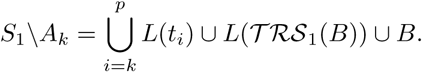

Thus, for any *k* ∈ [*w*(*e*_1_)], *π*_*k*_ is the bipartition induced by the *k*th edge on *P* (*e*_1_) in *T*_1_, where the edges are numbered from the side of *A*. Therefore, *π*_*k*_ ∈ *C*(*T*_1_) for any *k* ∈ [*w*(*e*_1_)]. Similarly, 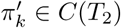 for any *k* ∈ [*w*(*e*_2_)].

Since for any *k* ∈ [*w*(*e*_1_)], *A*_*k*_ ∩*X* = *A* =*/ ∅* and (*S*_1_*\A*_*k*_)∩*X* = *B ≠ ∅*, we have *π*_*k*_|_*X*_ = *π* and *π*_*k*_ ∈ *Π*_*X*_. Similarly, for each *k* ∈ [*w*(*e*_2_)], *π*_*k*_^*t*^ ∈ *Π*_*X*_ and *π*_*k*_^*t*^ |_*X*_ = *π*. We also know that since *π ∉ C*(*T* |_*X*_), by Corollary 3, *π*_*k*_ *∉ C*(*T* |_*S*_1) for any *k* ∈ [*w*(*e*_1_)] and 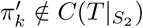 for any *k* ∈ [*w*(*e*_2_)]. We claim that 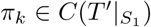 for all *k* ∈ [*w*(*e*_1_)] and 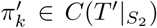 for all *k* ∈ [*w*(*e*_2_)]. Then assuming the claim is true, we have 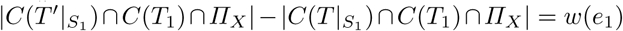 and 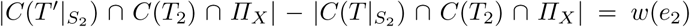, and thus *p*_*X*_ (*T ′*) *− p*_*X*_ (*T*) = *w*(*e*_1_) + *w*(*e*_2_) = *w*^*\**^(*π*).

Now we only need to prove the claim. Fix *k* ∈ [*w*(*e*_1_)], we will show that 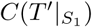. The claim of 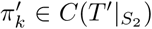 for any *k* ∈ [*w*(*e*_2_)] follows by symmetry. By invariant 2 of Lemma 10, we know that all extra subtrees of *𝒯 ℛ* (*e*_1_) were attached to *v* at the beginning of Algorithm 2 and thus the algorithm attaches them all onto (*v*_*a*_, *v*_*b*_) in the order of *t*_1_, *t*_2_, *…, t*_*p*_, such that *t*_1_ is closest to *A*. Let the attaching vertex of *t*_*i*_ onto (*v*_*a*_, *v*_*b*_) be *u*_*i*_ for any *i* ∈ [*w*(*e*_1_)]. Then we note *P* ((*v*_*a*_, *v*_*b*_)) is the path from *v*_*a*_ to *u*_1_, *u*_2_, *…, u*_*p*_ and then to *v*_*b*_. For any *t* ∈ 𝒯 ℛ 𝒮_1_(*A*), by invariant 3 of Lemma 10, *t* attached to *C*(*A*), the component containing *A* in *T ′*|_*X*_ *−* (*v*_*a*_, *v*_*b*_). Therefore, if we delete any edge of *P* ((*v*_*a*_, *v*_*b*_)) from *T ′, t* is in the same component as *A*. Similarly, for any *t* ∈ *𝒯 ℛS*_1_(*B*), *t* is in the same component as *B* if we delete any edge of *P* ((*v*_*a*_, *v*_*b*_)) from *T*. In particular, consider 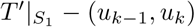. The component containing *u*_*k−*1_ and *A* contains all of *𝒯 ℛS*_1_(*A*) and {*t*_*i*_ | *i* ∈ [*k −* 1]}, thus the leaves of that component is

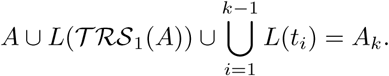

Therefore, the edge (*u*_*k−*1_, *u*_*k*_) induces the bipartition [*A*_*k*_|*S*_1_*\A*_*k*_] in 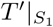. Hence, 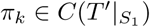 as desired. □

### Proposition 2

*Proof*. Let *weight*(*U*) denote the total weight of any set *U* of vertices of *G*. We first claim that *w*^*\**^(*I*) = *weight*(*I*^*\**^). Since all bipartitions in *C*(*T*_1_|_*X*_) ∩ *C*(*T*_2_|_*X*_) are compatible with all bipartitions in *C*(*T*_1_, *T*_2_, *X*), each of them become two isolated vertices in the weighted incompatibility graph, all of which are included in the maximum weight independent set *I*^*\**^. For each *π* ∈ *C*(*T*_1_|_*X*_) ∩ *C*(*T*_2_|_*X*_), the two vertices associated with it in *G* has total weight *w*(*e*_1_(*π*)) + *w*(*e*_2_(*π*)), which is exactly *w*^*\**^(*π*). For each *π* ∈ *C*(*T*_1_|_*X*_)*ΔC*(*T*_2_|_*X*_), the vertex associated with it also has weight exactly *w*^*\**^(*π*). Therefore, the *weight*(*I*^*\**^) = *w*^*\**^(*I*).

Fix any compatible subset *F* of *C*(*T*_1_, *T*_2_, *X*). Let *F ′* = *F \* (*C*(*T*_1 *X*_) ∩ *C*(*T*_2 *X*_)), and let *F ″* be a set of vertices constructed by combining the vertices associated with *F ′* and the two vertices associated with each of *π* ∈ *C*(*T*_1 *X*_) *C*(*T*_2 *X*_). Then *weight*(*F ″*) = *w*^*\**^(*F ′*) + *w*^*\**^(*C*(*T*_1_|_*X*_) ∩ *C*(*T*_2_|_*X*_)) *≥ w*^*\**^(*F ′*) + *w*^*\**^(*F ∩ C*(*T*_1_|_*X*_) ∩ *C*(*T*_2_|_*X*_)) = *w*^*\**^(*F*). Since *F* is compatible, *F ′* is also compatible, and thus *F ″* is an independent set in *G*. Therefore, *weight*(*F ″*) *≤ weight*(*I*^*\**^), since *I*^*\**^ is a maximum weight independent set in *G*. We conclude that *w*^*\**^(*F*) *≤ weight*(*F ″*) *≤ weight*(*I*^*\**^) = *w*^*\**^(*I*).□

We now present a lemma with the running time analysis for Algorithm 1, which complete the proof of Theorem 1.

### Lemma 11

*Algorithm 1 runs in O*(*n*^2^|*X*|) *time*.

*Proof*. First we analyze the running time of Algorithm 2, i.e., one refinement step. Dividing the neighbors of *v* and connecting them to *v*_*a*_ and *v*_*b*_ appropriately in line 3 *−* 6 take *O*(|*X*|^2^) time. We can do a depth-first-search in *T* |_*X*_ *− v* from every neighbor *u* of *v* and check in *O*(|*X*|) time if any newly discovered vertex is in *A* or *B* and connect *u* to *v*_*a*_ or *v*_*b*_ accordingly. Moving extra subtrees in *𝒯 ℛ\**(*π*) in line 7 takes *O*(*n*) time as *T*_*i*_ has at most *n* leafs and thus there are *O*(*n*) extra subtrees in total, so |*𝒯 ℛ* ^*\**^(*π*) | is *O*(*n*). Line 8 *−* 13 take *O*(*n*) time as the mappings are pre-calculated and there are again *O*(*n*) extra subtrees to be moved. Updating the data structures in line 15 *−* 21 takes *O*(|*X*| ^2^) time as there are at most *O*(|*X*|) bipartitions in *H*(*v*) and each of the containment conditions is checkable in *O*(|*X*|) time by checking whether one side of *π′* is a subset of one side of *π* (assuming that labels of leaves in both sides of the bipartitions are stored as pre-processed sorted lists instead of sets). The rest of the algorithm takes constant time. Overall, Algorithm 2 runs in *O*(*n* + |*X*|^2^) time.

Next we analyze the running time for Algorithm 1, i.e., Exact-RFS-2. Computing *C*(*T*_1_|_*X*_) and *C*(*T*_2_|_*X*_) in line 1 takes *O*(*n*^2^ + *n*|*X*|^2^) time as we need to compute *π*_*e*_|_*X*_ for all *e* ∈ *E*(*T*_1_) ∪ *E*(*T*_2_) and then take the union. There are *O*(*n*) edges in *E*(*T*_1_) ∪ *E*(*T*_2_). Computing *π*_*e*_|_*X*_ for each edge takes *O*(*n*) time by running DFS on *T*_*i*_*− e* to obtain *π*_*e*_ and then taking intersection of both sides of *π*_*e*_ with *X*, separately. Together it takes *O*(*n*^2^) time. Taking union of the bipartitions takes *O*(*n*|*X*|^2^) time as there are *O*(*n*) bipartitions to add and whenever we add a new bipartition, it needs to be compared to the *O*(|*X*|) distinct existing ones in the set. Since all bipartitions have size *O*(|*X*|), the comparison can be done in *O*(|*X*|) time (if each of them is represented by two sorted lists instead of two sets). In this step, we can always maintain a set of edges in *T*_*i*_ for each bipartition *π* ∈ *C*(*T*_1_, *T*_2_, *X*) such that *π*_*e*_|_*X*_ = *π*.

Line 2 *−* 5 compute the mappings and values we need in latter part of the algorithm. We analyze the running time for each *π* = [*A*|*B*] ∈ *C*(*T*_1_, *T*_2_, *X*) first. We can compute the path *P* (*e*_*i*_(*π*)) by assembling the set of edges associated with *π* in *T*_*i*_ from the last step into a path. This takes *O*(*n*^2^) time by counting the times any vertex appear as an end vertex in the set of edges. The two vertices appearing once are the end vertices of the path while those appearing twice are internal vertices of the path. Then *w*(*e*_*i*_(*π*)) = |*P* (*e*_*i*_(*π*))| can be found in constant time. Then we can find 𝒯 ℛ(*e*_*i*_(*π*)) by DFS in *T*_*i*_ *− v* for every internal node *v* of *P* (*e*_*i*_(*π*)), starting the search from the unique neighbor *u* of *v* such that *u* does not appear in the path. This takes *O*(*n*) time. We compute ℬ𝒫_*i*_(*A*) and ℬ𝒫_*i*_(*B*) by iterating over *O*(|*X*|) bipartitions in *C*(*T*_*i*_|_*X*_) and check if one side of any bipartition is a subset of *A* or *B* in *O*(|*X*|) time, this takes *O*(|*X*|^2^) time together. Next, we compute 𝒯 ℛ 𝒮_*i*_(*A*) (or 𝒯 ℛ 𝒮_*i*_(*B*)) by taking unions of extra subtrees in 𝒯 ℛ(*e*_*i*_(*π*)) for any *π* ∈ ℬ𝒫_*i*_(*A*) (or ℬ𝒫_*i*_(*B*)) in *O*(*n*) time. Extra subtrees are unique identified by their roots and 𝒯 ℛ(*e*_*i*_(*π*)) is disjoint from the set of extra subtrees associated with other edges, so taking union of at most *O*(*n*) extra subtrees takes *O*(*n*) time. Therefore, all the mappings and values can be computed in *O*(*n*^2^) time for each bipartition and thus it takes *O*(*n*^2^ |*X*|) time overall. With all the extra subtrees calculated for each partition, we can compute Extra(*T*_*i*_) in *O*(*n*^2^) time.

Constructing *T*_init_ in line 6 takes *O*(*n*) time. Line 7 constructs an incompatibility graph with *O*(|*X*|) vertices and *O*(|*X*|^2^) edges in *O*(|*X*|^3^) time as compatibility of any pair of bipartitions of size *O*(|*X*|) can be checked in *O*(|*X*|) time. For line 8, we can reduce Maximum Weight Independent Set to Minimum Cut problem in a directed graph with a dummy source and sink. Then the Minimum Cut problem can be solved by a standard Maximum Flow Algorithm. Since the best Maximum Flow algorithm runs in *O*(|*V* ||*E*|) time and the graph has *O*(|*X*|) vertices and *O*(|*X*|^2^) edges, this line runs in *O*(|*X*|^3^) time. Line 10-11 essentially runs line 7 of Algorithm 2 *O*(|*X*|) times using a total of *O*(*n*|*X*|) time. Line 12 initiates the data structure *H* and *sv* in *O*(|*X*|) time. Line 13 *−* 14 runs Algorithm 2 *O*(|*X*|) times with a total of *O*(*n*|*X*| + |*X*|^3^) time. Since |*X*| *≤ n*, |*X*|^3^ *≤ n*|*X*|^2^ *≤ n*^2^|*X*|, and thus, the overall running time of the algorithm is dominated by the running time of line 2 *−* 5, which is *O*(*n*^2^|*X*|).

We present additional results on the relationship between Relax–RFS and Relax–SFS and the hardness of RFS, SFS, and Relax–RFS.

### Lemma 12

*There exist instances of* Relax*–*RFS *and* Relax*–*SFS *in which an optimal solution to* Relax*–*RFS *is not an optimal solution to* Relax*–*SFS, *and vice-versa*.

*Proof*. Let *n ≥* 5 be any integer. Let *S*_*i*_ = [*n*] be the leaf set of *T*_*i*_ for all *i* ∈ [*n −* 3]. Let *π*_*i*_ = [1, 2, *…, i* + 1 | *i* + 2, *…, n*] for any *i* ∈ [*n −* 3]. We let *T*_*i*_ denote the tree with leaf set [*n*] that contains a single internal edge defining *π*_*i*_, and let *A* = {*T*_1_, *T*_2_, *…, T*_*n−*3_}. Let *T* be the star tree with leaf set [*n*] (i.e., *T* has no internal edges) and let *T ′* be the unique tree defined by *C*(*T*_*i*_) ⊆ *C*(*T ′*) for all *i* ∈ [*n*] (i.e., *T ′* is a compatibility supertree for *A*). Note that *T ′* is the caterpillar tree on 1, 2, *…, n* (i.e., *T ′* is formed by taking a path of length *n −* 2 with vertices *v*_2_, *v*_3_, *…, v*_*n−*1_ in that order, and making leaf 1 adjacent to *v*_2_, leaf *i* adjacent to *v*_*i*_, and leaf *n* adjacent to *v*_*n−*1_).

We will show that (1) *T* is an optimal solution for Relax–RFS(*A*), but not an optimal solution for Relax–SFS(*A*), and (2) that *T ′* is an optimal solution for Relax–SFS(), but not an optimal solution for Relax–RFS(*A*).

(1) We first show that *T* is not an optimal solution for Relax–SFS(*A*). Let *Π*_[*n*]_ denote the set of trivial bipartitions of [*n*]. Then *C*(*T*) = *Π*_[*n*]_. Let *Π′* = {*π*_*i*_ | *i* ∈ [*n −* 3]} (i.e., *Π′* contains the nontrivial bipartitions from the trees in profile *A*). Note that the set *Π′ ∪ Π*_[*n*]_ is compatible and that the caterpillar tree *T ′* (defined above) satisfies *C*(*T ′*) = *Π′ ∪ Π*_[*n*]_. Then *C*(*T ′*) ∩ *C*(*T*_*i*_) = *Π*_[*n*]_ ∪{*π*_*i*_} and thus SF(*T ′, T*_*i*_) = *n* + 1 for all *i* ∈ [*n −* 3]. Overall, the split support score of *T ′* is

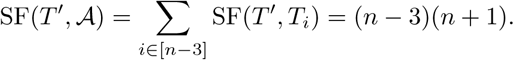

Since *C*(*T*) ∩ *C*(*T*_*i*_) = *Π*_[*n*]_, we have

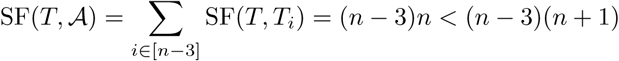

for any *n ≥* 5. Therefore, *T* is not an optimal solution for Relax–SFS(*A*).

Since |*C*(*T*)*\C*(*T*_*i*_)| + |*C*(*T*_*i*_)*\C*(*T*)| = 1 for all *i* ∈ [*n −* 3], the RFS score of T is

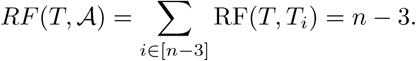

Now consider any tree *t ≠ T* with leaf set [*n*], and suppose *t* contains *p* bipartitions in *Π′* and *q* bipartitions in 2^[*n*]^*\*(*Π′ ∪ Π*_[*n*]_) where *p, q* ∈ N. Since *t ≠ T*, at least one of *p* and *q* is nonzero. Therefore,

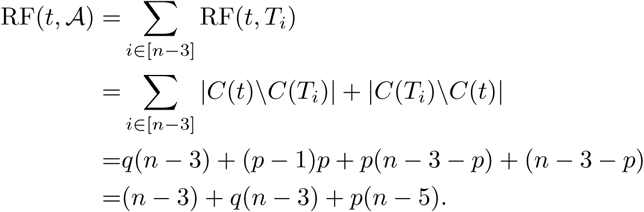

Since *n ≥* 5 and both *p* and *q* are non-negative with at least one of them nonzero, we know the RFS score of *t* is strictly greater than that of *T*. Therefore, *T* is an optimal solution to Relax–RFS(*A*).

For (2), the analysis above shows that *T ′* (since it is a compatibility supertree for *A*) has the largest possible split support score. Hence, *T ′* is an optimal solution to the relaxed Split Fit Supertree problem. However, the RFS score for *T ′* is (*n −* 4)(*n −* 3), which is strictly larger than *n −* 3 for *n >* 5, and the RFS score for the star tree *T* is *n −* 3; hence, *T ′* is not an optimal solution for the relaxed RF supertree problem.

We show that the Split Fit Supertree problem and the Asymmetric Median Supertree (AMS) problem, which was introduced in [46] and which we will present below, have the same set of optimal solutions and thus the hardness of one implies hardness of another. The input to the AMS problem is a profile 𝒜 = {*T*_*i*_ | *i* ∈ [*N*]} and the output is a binary tree

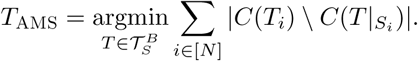

Thus, *T*_AMS_ minimizes the total number of bipartitions that are in the source trees and not in the supertree (i.e., *T*_*AMS*_ minimizes the total number of false negatives).

### Lemma 13

*Given profile 𝒜* = {*T*_1_, *T*_2_, *…, T*_*N*_} *and S* := _*i*∈[*N*]_ *L*(*T*_*i*_), *tree T* ∈ *𝒯*_*S*_ *is a Split Fit Supertree for 𝒜 iff T is an Asymmetric Median Supertree for 𝒜*.

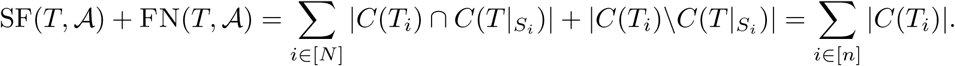

Hence, *T* is an Asymmetric Median supertree for 𝒜 iff *T* is a Split Fit supertree for 𝒜. □

### Lemma 6

RFS-3, SFS-3, *and* Relax*–*SFS-3 *are all* **NP***-hard*.

*Proof*. By Lemmas 1 and 13, for any profile *A*, the Robinson-Foulds, Split Fit, and Asymmetric Median supertree problems all have the same set of optimal solutions. Also, the Asymmetric Median Supertree problem is **NP**-hard for three trees when they have the same leaf set [30]; therefore, SFS-3 and RFS-3 are both **NP**-hard. Since refining a tree never decreases its split support score, SFS-3 trivially reduces to Relax–SFS-3, and thus Relax–SFS-3 is also **NP**-hard. □

## 9 Relationship between SMAST, SMCT, and RFS supertree problems

The SMAST and SMCT problems seek trees that are obtained after deleting minimal numbers of leaves from the input trees so that an agreement supertree or compatible supertree can be constructed from the reduced input trees. Here, we examine the possibility of using these output trees as constraint trees on the search for RFS supertrees, so that the removed taxa could be introduced into the constraint trees. We show that exact solutions to the SMAST and SMCT (Maximum Agreement Supertree and Maximum Compatible Supertree) problems are not directly relevent to solving the Robinson-Foulds supertree problem.

### Lemma 14

*There exists a pair of binary trees T*_1_ *and T*_2_ *for which some optimal SMAST or SMCT supertree cannot be extended to any optimal RFS supertree through the insertion of missing taxa*.

*Proof*. Consider the following two trees (both unrooted binary trees): Let *T*_1_ be given by the Newick string (*A*, ((*B, x*), ((*C, y*), (*D, E*)))) and let *T*_2_ be given by the Newick string (*A*, (*C*, (*z*, (*B*, (*D, E*))))).

An RFS supertree for this pair *T*_1_, *T*_2_ is given by (*A*, ((*C, y*), (*z*, ((*B, x*), (*D, E*))))), and has total RF distance to *T*_1_ and *T*_2_ equal to 2.

Note that at least one of *A, B, C* must be deleted to form an agreement supertree. Suppose *C* is deleted. Then ((*A, z*), ((*B, x*), (*y*, (*D, E*)))) is an optimal SMAST.

Observe that any way of adding *C* into this tree produces a supertree that has total RFS score greater than 2. Hence, for this pair *T*_1_ and *T*_2_ of input trees, for at least one optimal SMAST supertree, there is no way to extend that optimal supertree into an optimal RFS supertree.□

## 10 Maximum Weight Independent Set in Bipartite Graphs

Given an undirected bipartite graph *G*, with vertices *V* = *A ∪ B*, edges *E*, and vertex weights *w* : *V →* ℕ, the Maximum Weighted Independent Set problem tries to find a independent set *I ⊆ V* that maximizes *w*(*I*), where *w*(*S*) =∑ _*v S*_ *w*(*v*) for any *S ⊆ V*. It is well known (in folklore) that maximum weight independent set can be solved in polynomial time through reduction to the maximum flow problem. We reproduce a proof for completeness.

We first turn the graph into a directed flow network *G′* = (*V ∪{s, t*}, *E′*) where *s* and *t* are the newly added source and sink, respectively. To obtain *E′*, we direct all edges in *E* from *A* to *B*, add an edge from *s* to each vertex *u* ∈ *A* and add an edge from each vertex *v* ∈ *B* to *t*. We set the capacities *c* : *E′ →* N such that *c*(*e*) = *∞* if *e* ∈ *E, c*(*e*) = *w*(*u*) if *e* = (*s, u*) and *c*(*e*) = *w*(*v*) if *e* = (*v, t*). We claim that any *s, t*-cut (*S, T*) in *G′* has a finite capacity *k* if and only if (*S ∩ A*) ∪ (*T ∩ B*) is an independent set of weight *w*(*V*) *−k* in *G*.

We first observe that (*S ∩ A*) ∪ (*T ∩ B*) ∪ (*S ∩ B*) ∪ (*T ∩ A*) = (*S ∪ T*) ∩ (*A ∪ B*) = *A ∪ B* = *V*. Suppose (*S ∩ A*) ∪ (*T ∩ B*) is an independent set of weight *w*(*V*) *−k* in *G*. Since (*S ∩ A*) ∪ (*T ∩ B*) ∪ (*S ∩ B*) ∪ (*T ∩ A*) = *V*, the weight of (*S ∩ B*) ∪ (*T ∩ A*) is *w*(*V*) *−* (*w*(*V*) *−k*) = *k*. Since (*S ∩ A*) ∪ (*T ∩ B*) is an independent set, there is no edge from *S ∩ A* to *T ∩ B*. There is also no edge from *S ∩ B* to *T ∩ A* since edges in *E* are directed from *A* to *B*. Thus, the cut (*S, T*) consists of only edges from *s* to *T ∩ A* and from *S ∩ B* to *t*. Together the capacities of those edges equal the weight of the set (*S ∩ B*) ∪ (*T ∩ A*), which is *k*.

For the other direction of the proof, suppose (*S, T*) is an *s, t*-cut of finite capacity *k*. Since the cut has finite capacity, it does not contain any edge derived from *E*. In particular, there is no edge from *S ∩ A* to *T ∩ B* in *G′*, which implies there is no edge between *S ∩ A* and *T ∩ B* in *G*. Since there is also no edge among *S ∩ A* and *T ∩ B* in *G*, (*S ∩ A*) ∪ (*T ∩ B*) is an independent set. Since the edges in (*S, T*) solely consist of edges from *s* to *T ∩ A* and from *S ∩ B* to *t*, the sum of their capacities is *k*. Therefore, the weight of the set (*S ∩ B*) ∪ (*T ∩ A*) is *k* and the weight of (*S ∩ A*) ∪(*T ∩ B*) is *w*(*V*) *−k*.

Since *w*(*V*) is a fixed constant, we conclude that any *s, t*-cut (*S, T*) is a minimum cut in *G′* if and only if (*S ∩ A*) ∪ (*T ∩ B*) is an maximum weight independent set in *G*. By the standard Max-flow Min-cut theorem, a minimum *s, t*-cut in a directed graph is equivalent to the maximum *s, t*-flow. Thus, we can solve the Maximum Weighted Independent Set problem on bipartite graphs using a maximum flow algorithm in polynomial time.

## 11 Experimental study

We present additional details about the experimental performance study.

### 11.1 Datasets

*Experiment* 1 To produce the datasets for the experiment on multi-locus datasets, we used SimPhy [20] to generate species trees and gene trees with 501 species under the multi-species coalescent model, producing a set of true gene trees that differ from the true species tree by 68% of their branches on average due to ILS [19]. The number of genes varied from 25 to 1000 with ten replicate datasets per number of genes.

For each replicate dataset, we used the model species tree and a technique similar to DACTAL [26] (described below) to divide the species set into two overlapping subsets, each containing slightly more than half the species. ASTRAL [23, 24, 51] is a leading method for species tree estimation in the presence of ILS for large numbers of species, and so we used ASTRAL v5.6.3 (i.e., ASTRAL-III) to construct subset trees on the model gene trees, restricted to the relevant subset of species. Finally, the two ASTRAL subset trees were merged together using Exact-2-RFS and FastRFS. The following steps describe the procedure in details.

1. Identify a centroid edge (*a, b*) in the true species tree (i.e., an edge that, upon deletion, creates two subtrees *T*_*a*_ and *T*_*b*_ with leaf sets *A* and *B* of roughly equal size)
2. Let *X* be the set of 25 closest (in term of path distance on the weighted tree) leaves in *T*_*a*_ to *a* and in *T*_*b*_ to *b*
3. Let *A′* = *A ∪ X* and *B′* = *B ∪ X*
4. Restrict all 1000 true gene trees to leaf set *A′* and use ASTRAL-III [51] to compute a tree *A*1 on the restricted true gene trees
5. Restrict all 1000 true gene trees to leaf set *B′* and use ASTRAL-III to compute a tree *B*1 on the restricted true gene trees
6. Apply supertree methods FastRFS and Exact-2-RFS to input pair *A*1 and *B*1, and compare to the true species tree

We vary the number of true gene trees by selecting the first 100 and 25 true gene trees from the datasets with 1000 true gene trees.

*Experiment* 2 Each source tree is computed using maximum likelihood heuristics, with several clade-based source trees and a single scaffold source tree (i.e., species sampled randomly from across the tree). We selected the hardest of these 500-leaf conditions, where the scaffold tree has only 20% of the leaves. Because all the source trees miss some leaves, the number of leaves per supertree dataset varied. The source trees were then given to FastRFS and GreedyRFS to combine into a supertree.

We use the first 10 replicates out of a total of 30 replicates. Note that since replicate number 8 requires combining two trees with less than 2 shared taxa, supertree construction does not make sense on this replicate. After eliminating this replicate, we end up with 9 replicates in total. To make inputs with *k* source trees, for *k* ∈ {2, 4, 6, 8, 10, 12, 14}, we take the *first k* source trees in each replicate. Since the first tree is always the scaffold tree, all of our replicates contain the scaffold tree. The average number of leaves per each source tree per replicate for these datasets is (rounded to the nearest integer) {87, 79, 78, 73, 74, 73, 73}, for each corresponding *k*.

### 11.2 Scripts and Commands

Our scripts and other utilities (developed by the authors of this paper) are available at http://github.com/yuxilin51/GreedyRFS.

– *GreedyRFS on a set of source trees* `GreedyRFS.py -t <source_trees> -o <output_tree>` Note that when the input has two source trees, then GreedyRFS is identical to Exact-2-RFS.
– *RFS criterion score* To compute the RFS criterion score for a supertree *T* with respect to a profile *A*, we add the RF distances between *T* and every tree *t* ∈ *A*, as follows: `compare_trees.py <tree1> <tree2>`
– *Centroid decomposition (1 round)* `split_tree.py -t <input_tree> -o <output_directory>`
– *Find overlapping leaf set X* `find_x.py -t <input_tree> -o <output_directory>`
– *Restricting tree to a leaf set (Newick Utilities v1.6.0)* `nw_prune -v <input_tree> $(cat <label_of_leaves>) /` `> <output_tree>`

### 11.3 External software

*FastRFS v1.0*

`FastRFS -i <source_trees> -o <output_prefix>`

*SimPhy v1.0.2*

~~~
simphy -rs 10 -rl F:1000 -rg 1 -st F:500000 /
-si F:1 -sl F:500 -sb F:0.0000001 /
-sp F:200000 -hs LN:1.5,1 -hl LN:1.2,1 /
-hg LN:1.4,1 -su E:10000000 -so F:1/
-od 1 -v 3 -cs 293745 /
-o <output_directory>
~~~

*ASTRAL v5.6.3*

~~~
java -jar <path_to_astral_jar> -i <input_gene_trees> /
-o <output_est_species_tree>
~~~

